# Quantitative biophysical analysis of human septin hexamer and octamer self-assembly on model membranes

**DOI:** 10.64898/2026.06.01.729280

**Authors:** SaFyre Reese, Ward de Ridder, Annice van Hemmen, Alin-Gabriel Mateescu, Risa Togo, Shizue Omi-Massieau, Manos Mavrakis, Ralf P. Richter, Gijsje H. Koenderink

## Abstract

1

Septins are GTP-binding cytoskeletal proteins that shape and compartmentalize the plasma membrane. Their complex interactome has made it difficult to understand the molecular factors that govern their assembly. Moreover, it is unclear whether human septin hexamers and octamers form distinct higher-order assemblies, especially at the plasma membrane. Here, we address this question by using label-free methods to probe binding and self-assembly of recombinant human septins on supported lipid bilayers. Quartz crystal microbalance with dissipation (QCM-D) monitoring revealed that septin-membrane binding is diffusion-limited and concentration-dependent. Hexamers and octamers showed distinct viscoelastic properties, suggestive of structural differences. Imaging by atomic force microscopy (AFM) revealed that septin hexamers formed aligned nematic filamentous networks, whereas septin octamers formed aligned curved structures including spirals. QCM-D and AFM measurements both showed that septins form double-layered filament networks. However, upon C-terminal truncation of the SEPT6 and SEPT7 subunits, hexamers no longer bound the membrane while octamers formed a single-layered network of filament spirals. Our findings reveal that human septin hexamers and octamers interact differently with membranes, providing a baseline to understand their functions in the cell.

**Significance statement:**

- Septins are cytoskeletal proteins that control cell membrane shape and stiffness. It is poorly understood how septin oligomers, the basic building blocks of septin filaments, bind and assemble on membranes.
- We used label-free biophysical assays to quantitatively compare the binding kinetics and self-assembly behavior of recombinant human septin hexamers and octamers on supported lipid bilayer membranes.
- Our findings reveal that human septin hexamers and octamers both form organized filamentous networks on membranes, but with different structural properties that may potentially translate into different biological functions.

## 3 Introduction

Mammalian cells dramatically change their shapes as they divide, move, and shape tissues. They achieve this dynamic shape control by combining a flexible lipid bilayer membrane with a dynamic internal cytoskeleton. The mammalian cytoskeleton consists of four distinct protein filaments with complementary dynamic and mechanical properties. Actin filaments, microtubules, and intermediate filaments have well-studied roles in cell shape control (Conboy et al. (2024)). In contrast, the contribution of septins, the fourth component of the cytoskeleton, is less well understood (Schampera and Schwan (2024); Debadarshini et al. (2025)). However, there is growing evidence that they are essential for controlling the shape and mechanics of the cell surface during cellular shape changes. For instance, at the single-cell level, septins contribute to protrusion formation at the leading edge of crawling cells (Kho et al. (2023); Weiler et al. (2024)), uropod formation in motile T-cells (Gilden et al. (2012)), and membrane constriction and abscission of dividing cells (Ribet et al. (2017); Karasmanis et al. (2019); Carim and Hickson (2023)). They also modulate cell stiffness, either directly or through modulation of the actin cytoskeleton (Mostowy et al. (2011); Martins et al. (2023)). In a multicellular context, septins are important in collective cell migration (Gabbert et al. (2023)), maturation of neural tissues (Alkhanjari et al. (2025)), and tissue shape changes in developing embryos (Gabbert et al. (2023); Devitt et al. (2024)). Abnormalities in septin expression are linked with neurodegenerative and infectious diseases and are also hallmarks of cancer (Poüs et al. (2016); Robertin and Mostowy (2020); Alkhanjari et al. (2025)).

Septins are highly conserved in all eukaryotes except land plants (Delic et al. (2024)). They have been classified into different homology groups based on sequence homology. All septins share the same domain structure with a conserved GTP-binding domain (G-domain) flanked by highly variable N- and C-terminal domains. Pairs of septins from different homology groups form linear palindromic heterooligomers (Mendonça et al. (2019); Soroor et al. (2021)). Within these complexes, members of the same septin homology group can substitute for each other, a property referred to as the Kinoshita rule (Kinoshita (2003)). Within the cell, septins are found in the form of stable oligomers as well as filaments and higher-order networks. Biochemical reconstitution studies have shown that septin oligomers form linear filaments by end-to-end annealing, which in turn form higher-order structures such as bundles, networks and rings by interactions mediated by the long C-terminal coiled coil domains (reviewed in Cavini et al. (2021)).

*Homo sapiens* has 13 septin genes divided into 4 different homology groups named SEPT7 (SEPT7), SEPT3 (SEPT3-SEPT9-SEPT12), SEPT2 (SEPT1-SEPT2-SEPT4-SEPT5), and SEPT6 (SEPT6-SEPT8-SEPT10-SEPT11-SEPT14). Some septins are ubiquitously expressed across many tissues (e.g. SEPT2, SEPT6, SEPT7, SEPT9), while others (e.g. SEPT12 in testes and SEPT3 in brain tissue) are tissue-restricted. Further diversification occurs via alternative splicing and alternative promotors. At the genomic level, SEPT9 from the SEPT3 group is the most complex member of the septin gene family, coding for up to 18 isoforms (McIlhatton et al. (2001)). There are five main SEPT9 isoforms (SEPT9i1 to i5), which differ in the length and sequence of their N-terminus. These isoforms confer different functions on septin oligomers by tuning their interactions with microtubules, actin, organelles, and signaling molecules (Connolly et al. (2011); Targa et al. (2019); Devlin et al. (2021); Kuzmić et al. (2022); Okletey et al. (2023); Verdier-Pinard et al. (2017); Targa et al. (2019)). Three of the isoforms (SEPT9i1, i2 and i3) share a common long, disordered N-terminus. Of these, SEPT9i1 is frequently overexpressed in cancer and promotes cancer cell proliferation, survival and paclitaxel resistance owing to its interaction with microtubules (Targa et al. (2019); Kuzmić et al. (2022)) and stabilization of HIF1-*α* and JNK (Gonzalez et al. (2009); Amir et al. (2009)). Isolation of septins from human cells has revealed the existence of two sizes of stable heterooligomers: hexamers, composed of septins from the SEPT2, SEPT6, SEPT7 groups, and octamers containing additional septins (mostly SEPT9) from the SEPT3 group (Sellin et al. (2011); Kim et al. (2011); Sellin et al. (2014)). This contrasts with findings in other organisms, where septins form oligomers of a single size, such as octamers in *Saccharomyces cerevisiae* (budding yeast) (Frazier et al. (1998); Bertin et al. (2008); Bridges et al. (2014)) and hexamers in *Drosophila melanogaster* (fruit fly) (Field et al. (1996); Debadarshini et al. (2025)). While it is unclear whether human septin hexamers and octamers have distinct functions, interesting recent studies in an adherent human cell line showed that septin filaments that associate with actin stress fibers and microtubules contain exclusively octamers (Martins et al. (2023)). Moreover, there are many studies attributing specific biological functions to septin proteins that occur exclusively in octamers, such as SEPT9 or SEPT12 (Bai et al. (2016, 2013); Estey et al. (2010); Karasmanis et al. (2019); Kuo et al. (2015).)

Cell shape control by septins mainly depends on cortically localized septins. It is unclear whether the cortical septin pool in animal cells directly associates with the plasma membrane or indirectly associates with it via interactions with the membrane-bound actin cytoskeleton. Biochemical reconstitution studies suggest that both options are open, since mammalian septins are able to directly bind and polymerize on lipid membranes (Bertin et al. (2010); Bridges et al. (2014); Szuba et al. (2021); Nakazawa et al. (2023b)) but also bind actin filaments (Mavrakis et al. (2014); Smith et al. (2015); Iv et al. (2021)) and actin-binding proteins including anillin (Kinoshita et al. (2002)), non-muscle myosin-2 (Joo et al. (2007)), and the Cdc42 effector protein Cdc42EP3 (Tomasso et al. (2025)). In cells, septin and actin organization are both disrupted when either network is pharmacologically or genetically perturbed, suggesting an intimate reciprocal relationship between the two cytoskeletal systems (Kinoshita et al. (2002); Dolat et al. (2014); Martins et al. (2023)). The actin cytoskeleton forms distinct types of functional structures that co-localize with septins. In case of actin stress fibers, septins were recently shown to reside closer to the membrane than actin, suggesting that septins anchor actin stress fibers to the membrane (Martins et al. (2023)). In case of the actin cortex, septins have been identified as an important and likely filamentous component (Hagiwara et al. (2011); Park et al. (2015); Vadnjal et al. (2022)). However, the high density of the actin cortex has made it impossible to distinguish how septins are organized and whether they directly interact with the plasma membrane.

Biochemically reconstituting septins outside the complex environment of the cell has provided important insights in the intrinsic interactions of septins with membranes (reviewed in Woods and Gladfelter (2021); Nakazawa et al. (2023a)). One key question that has been explored with this approach is the lipid-selectivity of septins. Co-sedimentation assays of purified septins with liposomes revealed a specific affinity of mammalian septins for two types of phosphoinositide (PIP) lipids, PI(4,5)P_2_ and PI(3,4,5)P_3_ (Zhang et al. (1999)). Although PIP lipids are only a minority fraction (1-5%) of lipids in the cytoplasmic leaflet of the plasma membrane, they are essential signaling molecules and regulators of the cytoskeleton (Wills and Hammond (2022); Balla (2013)). Mitotic yeast septin octamers (capped by Cdc11) and human septin octamers have likewise been shown to selectively interact with PI(4,5)P_2_ lipids (Bertin et al. (2010); Nakazawa et al. (2023b)), although yeast septins are also able to bind phosphatidylinositol (PI) (Cannon et al. (2019); Vogt et al. (2025); Edelmaier et al. (2025)). Recent molecular docking analysis attribute the selectivity of Cdc11-capped yeast septin octamers for PI(4,5)P_2_ lipids to a binding pocket for the PI(4,5)P_2_ headgroup on the Cdc10 and Cdc11 subunits (Taveneau et al. (2020)). Yeast septin octamers capped by Shs1 (Taveneau et al. (2020)) and fly septin hexamers (Szuba et al. (2021)) were shown to lack PI(4,5)P_2_-selectivity, binding equally well to bilayers containing PI(4,5)P_2_ or phosphatidylserine (PS) lipids. When combined, PS and PI(4,5)P_2_ lipids synergistically enhance septin binding (Szuba et al. (2021); Beber et al. (2019)), suggesting that there is a substantial electrostatic component to septin-membrane binding. Indeed septin binding is disfavored at high (above 200 mM) monovalent ion concentrations (Szuba et al. (2021); Vogt et al. (2025); Goodchild et al. (2025)). The exact molecular determinants of membrane binding are not fully understood. Biochemical studies with purified monomeric septins suggest that positively charged polybasic domains (PB1 in the N-terminal domain and PB2 in the G-domain) bind anionic lipids Zhang et al. (1999); Casamayor and Snyder (2003); Bertin et al. (2010); Omrane et al. (2020), but the PB1 domains are likely buried in the interface between septin subunits (Castro et al. (2020); Cavini et al. (2021); Mendonça et al. (2021)). Additionally, conserved amphipathic helix domains have been identified at the C-terminus of the SEPT6 group (Cannon et al. (2019); Lobato-Márquez et al. (2021); Edelmaier et al. (2025)). Amphipathic helix domains insert into the membrane using lipid packing defects, which may contribute to septin’s ability to recognize membrane curvature (Bridges et al. (2016); Beber et al. (2019); El Alaoui et al. (2025); Goodchild et al. (2025); Curtis et al. (2026)).

Biochemical reconstitution has also been used to address a second important question, namely how membrane binding impacts membrane-templated septin self-assembly. Fluorescence imaging of yeast septins on solid-supported lipid bilayers showed that at low (3 nM) concentrations, septin octamers first associate with the membrane and then form filaments through 2D diffusion and end-on annealing (Bridges et al. (2014); Jiao et al. (2020)). Octamers and short filaments can grow into longer filaments via annealing or through cooperative recruitment of septin octamers from solution (Bridges et al. (2014); Jiao et al. (2020); Shi et al. (2023)). At higher (tens of nM) concentrations, yeast septin octamers form filamentous networks that are so dense that light microscopy cannot discern the individual filaments (Vogt et al. (2025); El Alaoui et al. (2025)). High-resolution imaging by electron microscopy (Bertin et al. (2010); Beber et al. (2019); Nakazawa et al. (2023b)) and atomic force microscopy (Szuba et al. (2021); Goodchild et al. (2025)) indicates that septins from various species (budding yeast, fly, and human septins) form dense networks of closely aligned filaments on membranes. Moreover, septin filaments form two-layered networks. In electron microscopy images, the two stacked layers appear to be orthogonal to each other with filaments in the first layer cross-bridged by filaments or oligomers in the second layer (Bertin et al. (2010); Beber et al. (2019); Szuba et al. (2021); Nakazawa et al. (2023b)). In atomic force microscopy images, alignment of the second layer was shown to be templated by the layer below (Goodchild et al. (2025)). It remains poorly understood how membrane-bound septin filaments are oriented relative to the surface. On bare mica and on electron microscopy grids (i.e., in absence of a lipid coating), septin filaments form train-track-like pairs with a 7 nm spacing which indicates that pairing is mediated by coiled coil oligomerization of the long and flexible C-termini (Bertin et al. (2010); Szuba et al. (2021); Jiao et al. (2020); Goodchild et al. (2025)). However, on supported lipid membranes, atomic force microscopy showed that filaments do not pair, suggesting that they may be re-oriented by 90 degrees, pointing their coiled coils up or down rather than sideways (Goodchild et al. (2025)). There is some evidence that the coiled coils may point downwards, allowing interactions of the C-terminal amphipathic helix domains of the SEPT6 subunits with the membrane (Jiao et al. (2020); Goodchild et al. (2025)). However, there is also some evidence that the coiled coils may point upwards facing the solution, because removal of the C-termini causes fly septin hexamers (Szuba et al. (2021)) and yeast septin octamers (Bertin et al. (2010)) to form monolayers, suggesting the coiled coils are exposed to the solution. Moreover, cryoelectron tomography of vesicle-bound human octamer septins showed that the first layer of septin filaments is in direct contact with the membrane, while the second layer is 12 nm apart from the first one, consistent with layer coupling via pairing of the coiled coil domains (Nakazawa et al. (2023b)).

So far the majority of biochemical reconstitution studies of septin-membrane binding have involved budding yeast septins. Several early studies used partially human septin hexamers, combining human SEPT6 and SEPT7 with mouse SEPT2 (Kinoshita et al. (2002); Mavrakis et al. (2014); Tanaka-Takiguchi et al. (2009)), or octamers, additionally containing human SEPT3 (DeRose et al. (2020)). Fully human recombinant septin oligomers are only available since 2021 (Iv et al. (2021); Castro-Linares et al. (2022); Fischer et al. (2022)). Human septin octamers have been shown to bind PI(4,5)P_2_-containing membranes (Martins et al. (2023)), forming two-layered networks that sense and generate micrometric membrane curvature (Nakazawa et al. (2023b)). However, the question whether septin hexamers and octamers, both present in cells, differ in their intrinsic biophysical properties remains open. Moreover, quantitative information on their binding kinetics are lacking.

Here we quantitatively compare the membrane binding kinetics and self-assembly behavior of purified human septin hexamers (composed of SEPT2, SEPT6 and SEPT7) and octamers (which additionally contain SEPT9i1). We measure the kinetics of septin binding to supported lipid bilayers using quartz crystal microbalance with dissipation monitoring (QCM-D), which measures septin (un)binding from the change in resonance frequency and dissipation of a membrane-coated quartz sensor. We furthermore image the septin networks using atomic force microscopy (AFM). Both methods show that human septin hexamers and octamers form two-layer filament networks in a two-step process, involving initial fast membrane recruitment of septins followed by a slower process of ongoing septin recruitment and network remodeling. We also show that hexamers and octamers form differently organized filament networks and reveal a key role for the SEPT6 and SEPT7 coiled coils in membrane binding and septin self-assembly, respectively. Our findings demonstrate that, in isolation, human septin hexamers and octamers exhibit different interactions and self-assembly properties on membranes.

## 4 Results

### 4.1 Human septins form rigid membrane-bound films with a self-limiting thickness

To measure the kinetics of septin binding to lipid membranes, we employed quartz crystal microbalance with dissipation monitoring (QCM-D). This acoustic technique allows time-resolved and label-free measurements of protein binding to the surface of the quartz crystal sensor with high (1 s) time resolution. Protein binding is detected by the decrease in the resonant frequency of an oscillating piezoelectric sensor as proteins adsorb to the sensor and add mass. The frequency shift (Δ*F* ) is proportional to the amount of protein, plus the hydrodynamically trapped water in the network, that is bound to the sensor surface (Reviakine et al. (2011)). Simultaneously, the dissipation (Δ*D*), measured as energy lost per oscillation cycle, describes the mechanical properties of the material, which dampens sensor oscillations (Johannsmann et al. (2009)). The first step of each experiment was to form a PS- and PI(4,5)P_2_-containing supported lipid bilayer (SLB) on the QCM-D sensor by perfusing the channel with a suspension of small unilamellar vesicles (SUVs) (see Figure 1A). After injection of SUVs, Δ*F* (blue line) decreased as the intact SUVs bind and add mass to the sensor; simultaneously, Δ*D* (red line) increased as more energy was dissipated, due to the softness of the SUVs (step 2 in Figure 1B). Once the surface is covered, the SUVs rupture and coalescence into an SLB (Richter and Brisson (2005)). The final values of Δ*F* ( ≈ 28 Hz) and Δ*D* (below 0.5 × 10^-6^) confirm the formation of an thin, rigid lipid bilayer (Richter et al. (2006)).

**Figure 1:**
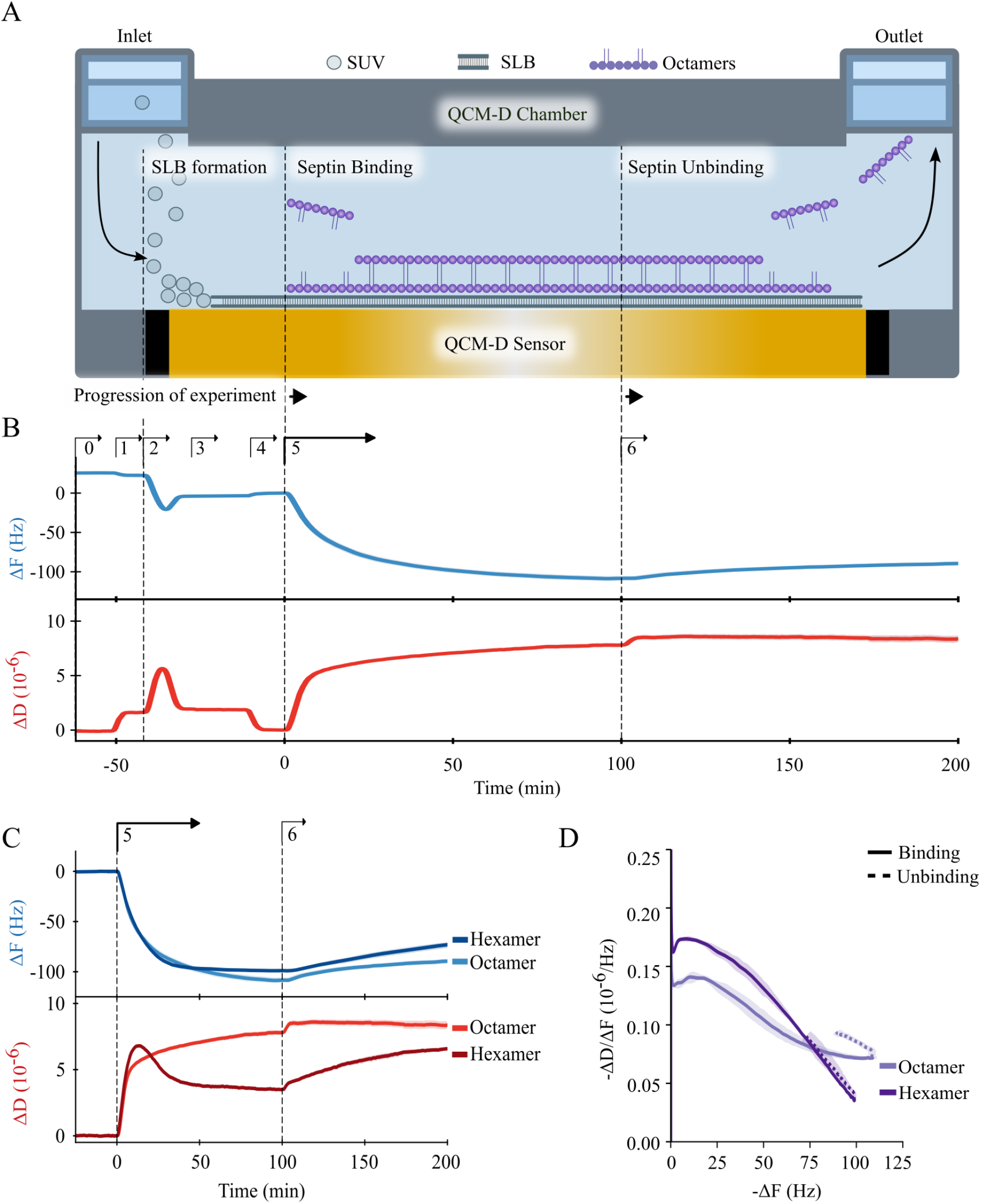
QCM-D measurements of septin binding to supported lipid bilayers (SLBs). (A) Schematic overview of the QCM-D sensor and the workflow of the experiments. *Left*: first, the channel is perfused with small unilamellar vesicles (SUVs) to form an SLB on the sensor. *Middle*: next, septin oligomers are perfused into the system. *Right*: once septin-membrane binding has equilibrated, buffer is perfused to investigate the reversibility of binding. (B) Example QCM-D experiment with septin octamers (50 nM) from start to finish, with a constant flow rate of 20 µL min^−1^. Top panel shows frequency shift, bottom panel shows corresponding dissipation shift. Arrows with numbers indicate sequential perfusion of: (0) low-salt polymerization buffer to set baseline; (1) exchange to citrate buffer; (2) SUVs in citrate buffer; (3) citrate buffer to remove remaining SUVs; (4) exchange to low-salt polymerization buffer; (5) septin octamers in low-salt polymerization buffer; (6) low-salt polymerization buffer to check reversibility of binding. C) Membrane binding of (5) hexamers (dark curves, *n* = 2) and octamers (light curves, *n* = 2) (both at 50 nM), followed by (6) partial desorption in low-salt polymerization buffer. Curves show data variation. Note that only steps 5 and 6 of the full experiments illustrated in panel C are shown. (D) Softness parameter ( − Δ*D/*Δ*F* ) (calculated from data in panel (C) as a function of − Δ*F*, which is a measure for surface coverage. Solid lines represent binding phase (step 5) while dashed lines represent unbinding phase (step 6). Lines represent the mean and shaded areas the data spread. For buffer compositions: see Table 1.

After perfusing the lipid bilayer with buffer to remove any remaining SUVs (steps 3 and 4), we injected purified human septin oligomers (step 5 in Figure 1B). This produced an immediate sharp decrease in Δ*F* and a concurrent increase in Δ*D*, both for octamers and hexamers (Figure 1C). These rapid changes indicate that both oligomer species rapidly bind the membrane. After about 100 minutes, the frequency shifts reached a plateau despite continued perfusion with septin oligomers, indicating the presence of a mechanism that limits septin binding. The steady state values of Δ*F* were comparable for hexamers ( ≈ −90 Hz) and octamers ( ≈ −110 Hz), indicating that they form protein films with comparable thickness. For rigid and spatially uniform films, we can estimate this thickness from the measured frequency shift using a modified Sauerbrey equation (Sauerbrey (1959); Richter et al. (2003); Reviakine et al. (2011)). The septin films are sufficiently rigid to justify this approximation, given that their softness — defined as the ratio of Δ*D*/-Δ*F* — is well below the threshold value of 0.4 × 10^−6^ Hz^−1^ for the Sauerbrey equation to hold (Figure 1D) (Reviakine et al. (2011)). According to the Sauerbrey equation, the frequency shift is proportional to the product of the mass density of the protein film (*ρ*) and its thickness (*d*):

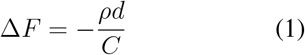

Here, Δ*F* is the normalized frequency shift (see Methods section 6.6) and *C* is the mass sensitivity constant of the QCM-D sensor (18 ng cm^−2^ Hz^−1^ (Reviakine et al. (2011)). The mass density (*ρ*) of the protein film should be somewhere in-between the mass density of the solvent (1 g cm^−3^) and that of the protein (typically 1.4 g cm^−3^ (Fischer et al. (2004)). Assuming a septin volume fraction of 10%, we obtain a mass density *ρ* of 1.04 g cm^−3^ (see Methods section 6.6 for details). From the frequency shifts measured after 100 min, we estimate that hexamers and octamers indeed reach comparable film thicknesses of 16.9 ± 0.5 nm (*n* = 4) and 17.2 ± 1.2 nm (*n* = 7), respectively. Note that, since we set the frequency shift to zero after SLB formation, this film thickness corresponds to the thickness of the septin film only. Importantly, the film height estimation is rather insensitive to the septin volume fraction. If we, for instance, assume a septin volume fraction of 50%, the film thicknesses are 14.6 ± 1.4 nm and 14.9 ± 1.7 nm, respectively. These values exceed the dimensions of septin monomers, which are defined by the size of their globular domain (4 nm) plus some additional contribution by the C-terminal coiled coils of the SEPT2, SEPT6 and SEPT7 subunits (Sirajuddin et al. (2007)). The estimated contour lengths of the coiled coils are 4 nm for SEPT2 and 13 nm for SEPT6 and SEPT7 (Szuba et al. (2021)). Moreover, they are flexible and connected to the G-domain by unstructured flexible hinges (Bertin et al. (2008); Sirajuddin et al. (2007); Szuba et al. (2021)). Recent AFM experiments of yeast septin filaments on membranes suggest they protrude from the bilayer surface by only 4.9 nm (Goodchild et al. (2025)). We thus consider it unlikely that the 17 nm film thickness represents a septin monolayer. The septin film thickness is comparable to the 19 nm thickness measured previously for fly septin hexamers on SLBs (Szuba et al. (2021)) and is also close to the predicted 20 nm spacing between septin filaments connected by antiparallel SEPT6 and SEPT7 coiled coils (Leonardo et al. (2021)). Indeed, several studies of septin-membrane binding (fly, yeast, human) have shown by AFM or electron microscopy that septins form double-layered networks (Beber et al. (2019); Szuba et al. (2021); Nakazawa et al. (2023b); Goodchild et al. (2025)). Based on the QCMD data we therefore hypothesize that human septin hexamers and octamers form two-layered networks in the QCM-D experiments, a hypothesis we will revisit below by AFM imaging.

Interestingly, we find that septin binding is accompanied by qualitatively different time dependencies of the dissipation shifts for human septin hexamers and octamers (Figure 1C bottom). At early times, the dissipation shift rapidly increases in both cases, indicative of rapid recruitment of septin oligomers from the solution to the membrane. For hexamers, the dissipation shift shows a transient peak at intermediate coverage (reached around t = 13 minutes) followed by a decrease indicating an apparent mechanical stabilization of the adsorbed layer with increasing surface coverage. Similar transient dissipation peaks have been observed previously for globular proteins adsorbing to membranes (Höök et al. (2001); Richter and Brisson (2005); Bingen et al. (2008)). Computer simulations modeling proteins as spheres tethered to the membrane by a flexible anchor showed that the dissipation reduction results from hydrodynamic coupling of the proteins to the shear deformation applied by the oscillating QCM-D sensor (Johannsmann et al. (2009)). At low surface coverage, the dissipation is large because the particles are able to rock around their flexible attachment in the flow field of the background fluid. At high surface coverage, lateral hydrodynamic interactions between neighboring particles counteract this mode of dissipation, resulting in a maximum in dissipation at intermediate adsorption times. The observation of a dissipation peak for septin hexamers is therefore indicative of the presence of flexible linkers in the contact zone between the proteins and the surface. Interestingly, we do not observe any dissipation peak for septin octamers, which instead show a monotonic increase of the dissipation shift, before reaching a limiting value (Figure 1C bottom, lighter curves). The dissipation responses suggest that septin hexamers and octamers bind similarly at early stages, but that the mechanics of the film are different in later stages.

To test the reversibility of septin binding, we perfused low-salt (50 mM KCl) polymerization buffer once binding had reached steady state (step 6 in Figure 1B). These buffer conditions should maintain septins in a polymeric state (Iv et al. (2021)). In case septin-membrane binding is reversible we expect that incubation with buffer should cause septin desorption to equilibrate the membrane-bound and solution pools. As shown in Figure 1C, there was indeed a gradual loss of protein upon buffer incubation, but this loss was only partial. Over a period of 100 minutes, the total fraction of desorbed protein (expressed in terms of the fractional change in frequency shift) was 25.6% ± 2.8% for hexamers and 17.5% ± 0.04% for octamers. The fact that a major fraction of protein remained bound indicates that septins rather stably adsorb to the membrane. When we perfused the channel with high-salt (300 mM KCl) buffer to induce septin depolymerization (Iv et al. (2021)), we observed additional detachment of 75.9% ± 0.4% (*n* = 2) for the septin hexamers and 30.2 % ± 0.8% (*n* = 2) for the human octamers (see Supplementary Figure 1A).Septin desorption in high-salt buffer could be an effect of septin depolymerization and/or electrostatic interactions (Szuba et al. (2021); Vogt et al. (2025)). When we finally perfused the QCM-D channel with guanidinium chloride and buffer to denature and remove remaining membrane-bound septins, the frequency and dissipation values returned to the bare membrane values measured before incubation with septins (Supplementary Figure 1A). This observation shows that septin binding does not perturb the lipid bilayer, so membrane binding is likely peripheral (Arwin (1998)).

To investigate how the mechanical properties of the septin films evolve during the binding process, we calculated the softness parameter, defined as the ratio of the dissipation change over the frequency shift, Δ*D*/-Δ*F* . In previous QCM-D studies of globular proteins, this parameter was shown to provide a measure of elastic compliance due to flexible hinges linking the proteins to the surface (Johannsmann et al. (2009); Reviakine et al. (2011)). To facilitate a side-by-side comparison of septin hexamer and octamer films, we plot the softness as a function of -Δ*F*, a quantity that is proportional to the adsorbed areal mass density or surface coverage (solid lines in Figure 1D). For both oligomer species, the softness starts to decrease monotonically with surface coverage for values of -Δ*F*) above 25 Hz, suggesting that septins form an increasingly rigid film once the coverage exceeds a certain threshold. However, the surface coverage dependence of the softness is steeper for the septin hexamers than for the octamers, culminating in a smaller final softness for hexamers (0.03 × 10^−6^ Hz^−1^) as compared to octamers (0.07 × 10^−6^ Hz^−1^). Hexamers thus form more rigid films than octamers, suggesting differences in spatial organization of the septin films. The softness values are comparable to those of membrane-bound fly septin hexamers (0.07 × 10^−6^ Hz^−1^) (Szuba et al. (2021)). Septin films are significantly less rigid than rigidly attached streptavidin monolayers ( ≈ 0.015 × 10^−6^ Hz^−1^) and closer in rigidity to streptavidin monolayers tethered by flexible DNA linkers ( ≈ 0.1 × 10^−6^ Hz^−1^) (Johannsmann et al. (2009)). This suggests that the septin films contain flexible molecular linkers. Likely candidates are the C-terminal coiled coils or their flexible linkages to the globular domains of the SEPT2, SEPT6 or SEPT7 subunits or the flexible N-terminus of SEPT9. Upon perfusion with low-salt buffer, partial septin desorption (visible from a decrease of the frequency shift) was accompanied by an increase of the dissipation shift (dashed lines in Figure 1D). For septin hexamers, the softness parameter during septin desorption increased back along the same (but reversed) trajectory followed during septin binding. In contrast, the softness parameter during octamer unbinding increased along a different trajectory of consistently higher softness than during the octamer binding process. This hysteresis suggests that the octamer films undergo irreversible structural changes during binding.

### 4.2 Septin-membrane binding is mass-transport limited

The rate of protein adsorption on membranes is in general expected to be limited by either transport (via diffusion and flow) to the surface or the reaction-limited rate of attachment to the membrane (Hermens et al. (2004)). To test which process limits the initial rate of septin-membrane binding, we performed QCM-D measurements of septin-membrane binding (at 50 nM septin concentration) for a range of flow rates (*Q*) between 5 µL min^−1^ and 100 µL min^−1^. In case of mass-transport-limited binding, the binding rate should go up with increasing flow rate. By contrast, if membrane binding itself is rate-limiting, the binding rate should be independent of flow rate. The time traces of the frequency and dissipation shifts show a clear increase of binding rate with increasing flow rate for human septin hexamers (Figure 2A) as well as octamers (Figure 2B), demonstrating that binding is diffusion-limited. To quantify the flow rate dependence, we extracted the maximal binding rate from the first derivatives of the time traces for each individual experiment and plotted the binding rates as a function of flow rate on a double-logarithmic scale. This revealed straight lines indicative of a power law functional dependency with exponents of *α* = 0.62 ± 0.10 (95% CI) for septin hexamers (Figure 2C) and *α* = 0.53 ± 0.14 (95% CI) for septin octamers (Figure 2D). This scaling falls in-between theoretical limits of 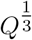-scaling and *Q*^1^-scaling predicted for protein adsorption within a straight channel under constant laminar flow. Convective fluid transport leads to the formation of a surface-proximal layer that is depleted in proteins. The weaker 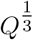 -scaling is expected when this depletion layer is thinner than the height of the flow channel (Hermens et al. (2004)), whereas the stronger *Q*^1^-scaling is expected when the solution above the surface becomes depleted of proteins throughout the full channel height (Kirichuk et al. (2023)). The intermediate scaling exponents we observe indicate that mass transport is limited by a layer depleted of septins above the membrane.

**Figure 2:**
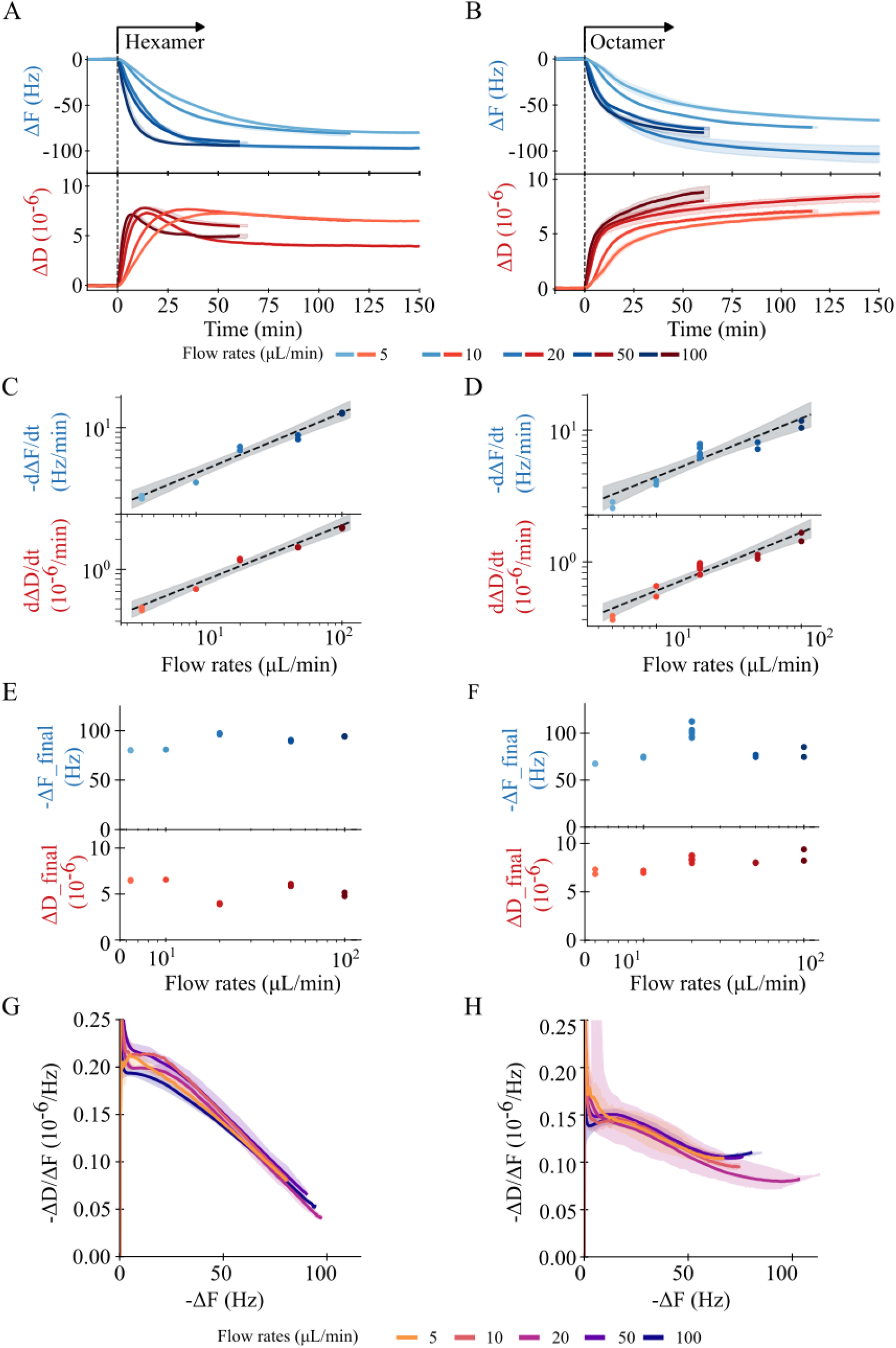
QCM-D experiments at different flow rates show that septin-membrane binding is initially mass-transport limited. (A) Septin hexamer (50 nM) binding at different flow rates (see legend at bottom), showing the frequency shift (top) and dissipation shift (bottom). Arrow indicates start of hexamer perfusion. Shaded areas represent the total spread of the data. (B) Same for septin octamers (50 nM). (C) Flow rate dependence of the maximal binding rates, for frequency shift (top) and dissipation shift (bottom) for hexamers. The black dashed line is a linear fit to the log-transformed data with exponent 0.62 ±0.10, with the shaded area representing the 95% confidence interval of the fit. (D) Same for octamers, with best-fit power-law exponent of 0.53 ±0.14. (E) Steady-state frequency shifts (top) and dissipation shifts (bottom) for hexamers. (F) Similar data for octamers. (G) Softness parameter (− Δ*D/*Δ*F* ) for septin hexamers as a function of − Δ*F* . H) Similar data for octamers. Shaded areas represent the total spread of the softness parameter for each flow rate dataset. Legends at the bottom explain the color code for flow rates in panels A-F (left) and panels G-H (right). Flow rates for hexamers were 5 µL min^−1^ (*n* 2), 10 µL min^−1^ (*n* = 1), 20 µL min^−1^ (*n* = 2), 50 µL min^−1^ (*n* = 2), 100 µL min^−1^ (*n* = 2). Flow rates for the octamers were 5 µL min^−1^ (*n* = 2), 10 µL min^−1^ (*n* = 1), 20 µL min^−1^ (*n* = 7), 50 µL min^−1^ (*n* = 2), 100 µL min^−1^ (*n* = 2).

In contrast to the marked dependence of binding rate of septins on flow rate, the final frequency and dissipation shifts did not show any systematic dependence on the flow rate, neither for hexamers (Figure 2E) nor for octamers (Figure 2F). Also, the curves of softness (Δ*D*/-Δ*F* ) versus surface coverage (-Δ*F* ) collapsed onto a single trajectory irrespective of flow rate for both the hexamers (Figure 2G) and octamers (Figure 2H). Together, these observations suggest that the surface density and structure of the resulting septin films are not significantly impacted by the presence of flow.

### 4.3 Concentration dependence of septin-membrane binding

To test the impact of septin concentration on membrane binding, we performed a series of QCM-D measurements with septin oligomer concentrations between 6.25 nM and 50 nM. Both for hexamers (Figure 3A) and octamers (Figure 3B), the QCM-D time traces revealed a linear relationship between increased concentration and increased initial rate of binding, evident from a steeper change of the frequency shifts (blue lines) and dissipation shifts (red lines) (Figure 3C-D). At the same time, also the final amount of bound septins after 150 minutes increased with septin concentration, as evidenced by increasing final frequency and dissipation shifts for hexamers (Figure 3E) and octamers (Figure 3F).

**Figure 3:**
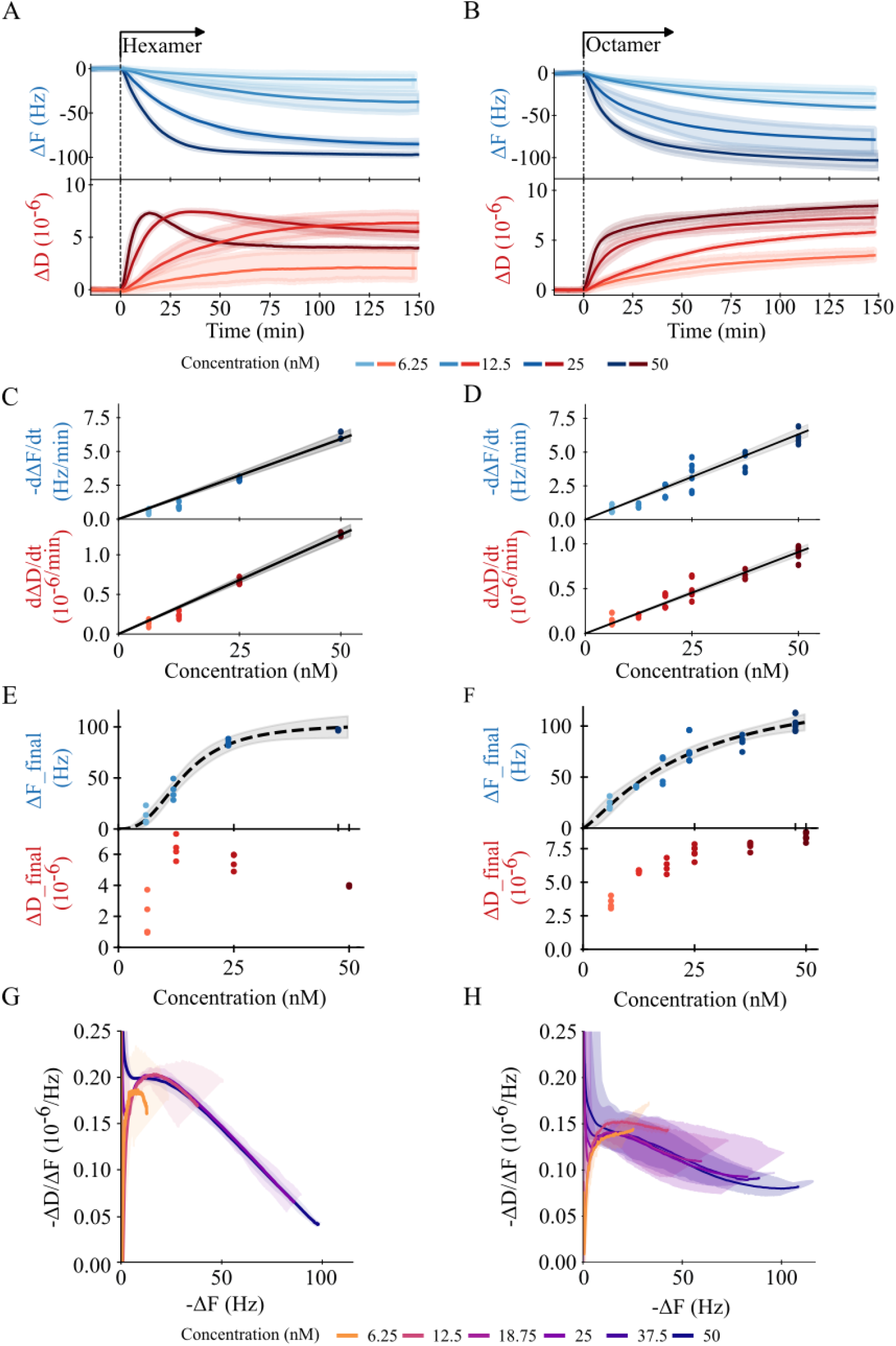
QCM-D measurements show that septin binding to SLB membranes is concentration dependent but the viscoelasticity of the septin films is largely independent of concentration. (A) Time-dependent frequency shift (top) and dissipation shifts (bottom) for human septin hexamers at different concentrations between 6.25 nM and 50 nM (see legend at bottom) using a constant flow rate of 20 µL min^−1^. (B) Same for septin octamers. The shading represents the total spread of the data. (C) Concentration dependence of the maximal binding rate for hexamers, from Δ*F* (top) and Δ*D* (bottom), obtained from panel A. The black lines are linear fits, constrained to go through the origin, with slopes 0.118 ± 0.009 Hz min^−1^ nM^−1^ (top) and 0.025 × 10^−6^ ± 0.0013 × 10^−6^ min^−1^ nM^−1^ (bottom). (D) Same for octamers. The slopes of the fits are 0.126 ± 0.009 Hz min^−1^ nM^−1^ (top) and 0.018 × 10^−6^ ± 0.0009 × 10^−6^ min^−1^ nM^−1^ (bottom). (E) Concentration dependence of the steady-state frequency shifts (top) and dissipation shifts (bottom) for hexamers. The dashed black line is the Hill equation fit with *n*_H_ = 2.68 ± 0.94 and *K* = 14.8 ± 2.7 nM. (F) Similar data for octamers, with fitted *n*_*H*_ = 1.24 ± 0.65 and *K* = 24.8 ± 22.1 nM. (G) Softness parameter ( − Δ*D/*Δ*F* ) for hexamers. H) Softness parameter for octamers. The shaded area represents the total spread of the data for panels A, B, G, and H. The reported margin or error of the fit slope and shaded area represent the 95% CI of the fit for panels C-F. The hexamer concentrations (during the binding phase) were 6.25 nM (*n* = 4), 12.5 nM (*n* = 4), 18.75 nM (*n* = 4), 25 nM (*n* = 4), 37.5 nM (*n* = 4), 50 nM (*n* = 2). The octamer concentrationswere 6.25 nM (*n* = 4), 12.5 nM (*n* = 4), 18.75 nM (*n* = 4), 25 nM (*n* = 6), 37.5 nM (*n* = 4), 50 nM (*n* = 7).

Since the effective binding isotherms showed sigmoidal dependencies on septin oligomer concentration, we fitted them with Hill functions (see Methods 6.6). The fitted effective Hill coefficients were 2.68 ± 0.94 for hexamers and 1.24 ±0.65 for septin octamers, indicating that septin oligomers engage in cooperative membrane binding, likely as a consequence of septin polymerization. Note that these effective Hill coefficients are expected to represent lower limits on the real Hill coefficients because the contribution of hydrodynamically trapped solvent to the frequency shifts is typically found to decrease with protein coverage (Reviakine et al. (2011)). The different degrees of binding cooperativity for hexamers and octamers suggests a mechanistic difference between septin network organization and membrane binding between septin hexamers and octamers.

To quantify the concentration dependence of the binding rate, we determined the initial septin binding rate from the frequency time traces. We used the Sauerbrey equation to convert the frequency shift time derivatives into the septin mass binding rate (for details see Methods section 6.6). The initial septin binding rate increased linearly with the bulk septin concentration with a pseudo-first order rate constant of 0.075 ± 0.006 s^−1^ µm^−2^ nM^−1^ for hexamers (Figure 3C) and 0.055 ± 0.004 s^−1^ µm^−2^ nM^−1^ for octamers (Figure 3D). These binding rates are about an order of magnitude lower than those reported for binding of yeast septin octamers to spherical SLB-coated beads, which were of order 1 s^−1^µm^−2^nM^−1^ measured using single molecule fluorescence microscopy (Shi et al. (2023)). However, we note that these rates are effective rates since septin-membrane binding is mass transport-limited.

The dissipation shift time traces show an interesting dependence on septin concentration. For hexamers, we observe a sigmoidal increase of the dissipation shift at the two lowest hexamer concentrations (6.25 nM and 12.5 nM), whereas the dissipation shift displays transient peaks at the two higher hexamer concentrations (25 nM and 50 nM) (Figure 3A).

This behavior indicates that at low hexamer concentrations the surface coverage is too low to allow for lateral hydrodynamic interactions between neighboring proteins, whereas at higher hexamer concentrations the surface coverage is high enough to suppress dissipation by hydrodynamic interactions. The final dissipation shift reached at steady state strongly increases when the septin hexamer concentration is increased from 6.25 nM to 12.5 nM, but decreases somewhat with increasing concentration above 12.5 nM, consistent with the suppression of dissipation by increased surface crowding (Figure 3E). For octamers, we instead observe sigmoidal time traces for the dissipation shift at all concentrations, without any transient peaks (Figure 3B). Moreover, the final dissipation shift monotonically increases with septin octamer concentration over the entire concentration range (Figure 3F). This observation suggests the presence of structural differences of some kind between hexamer and octamer septin films. The softness (-Δ*D*/Δ*F* ) decreases with surface coverage (-Δ*F* ) along the same trajectory irrespective of septin concentration, except at the lowest concentrations, 6.25 nM for hexamers (Figure 3G) and 6.25 nM and 12.5 nM for octamers (Figure 3H). The collapse of the curves at the higher septin concentrations suggests the process of septin binding and self-assembly is independent of septin concentration and binding rate at higher surface coverage, as already observed earlier when modulating septin binding by the flow rate (Figure 2G-H)

To determine the characteristic time scales associated with the septin-membrane binding process, we performed an alternative phenomenological analysis of the time traces for the frequency and dissipation shifts, fitting them to (multi-)exponential curves (Supplementary Figure 2A and B). At low septin concentrations (6.25 nM and 12.5 nM), the binding curves were well described by a single exponential function for hexamers (Figure 4A) as well as octamers (Figure 4B), suggesting that the binding process involves a single process captured by a single timescale. In contrast, at higher septin concentrations (25 nM or 50 nM), we needed double-exponential fits to capture the time traces for hexamers (Figure 4C) and octamers (Figure 4D), suggesting that the binding process involves a fast and a slow process. A triple-exponential fit showed that the single- and double-exponential fits were sufficient to capture the behavior of the septin binding (Supplementary Figure 2). The short and long timescales both decreased with increasing septin concentration, for hexamers (Figure 4E) as well as octamers ((Figure 4F). Overall, the binding rate was slightly higher for hexamers than for octamers.

**Figure 4:**
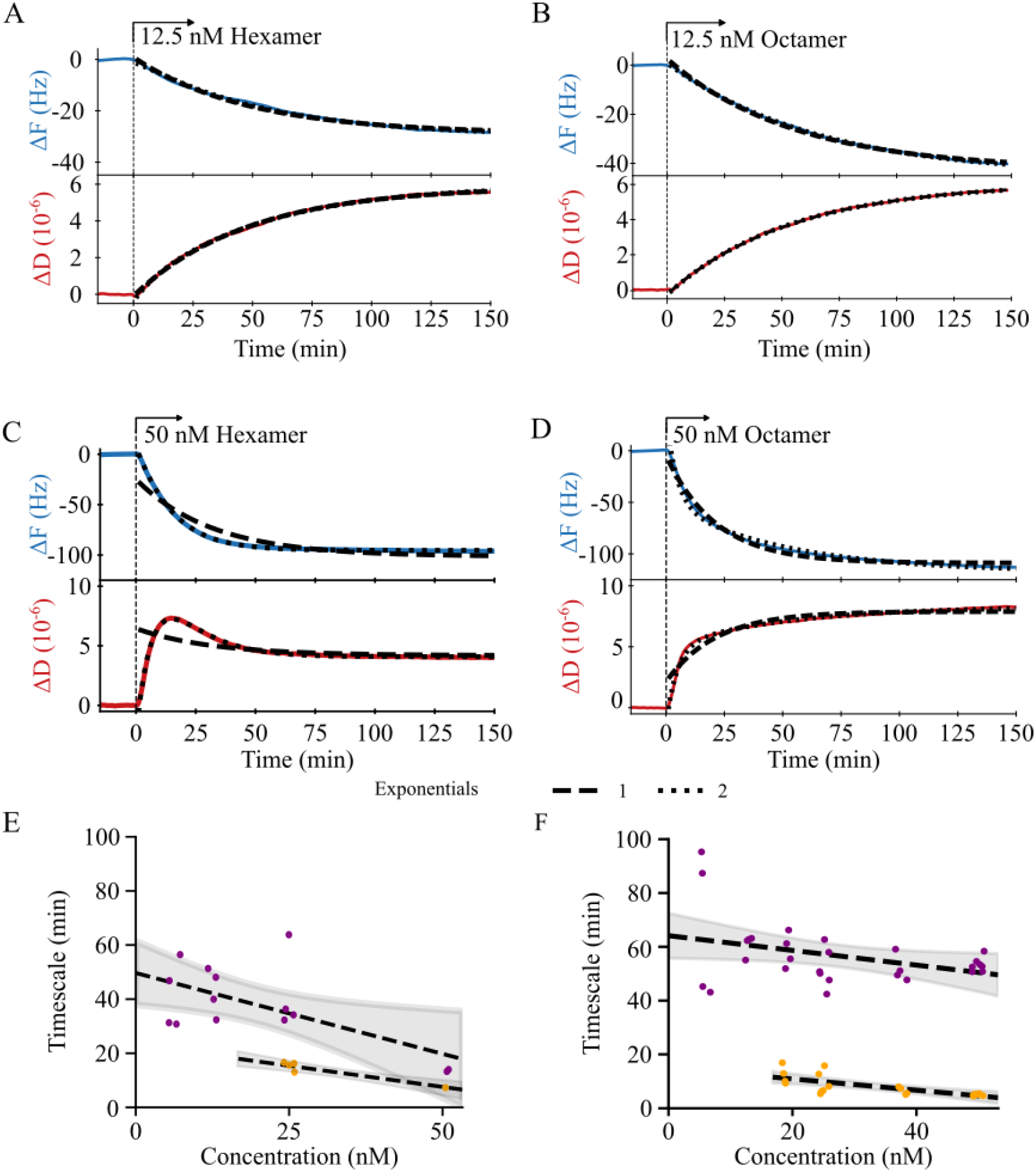
QCM-D measurements show that human septin hexamers and octamers bind SLB membranes with two distinct timescales. (A) QCM-D time traces for 12.5 nM septin hexamers (*n* = 4) are well-described by a single-exponential fit (long-dashed line). (B) Same for 12.5 nM octamers (*n* = 4). (C) Time traces for 50 nM hexamers (*n* = 2) are not captured by a single-exponential fit (long-dashed line) but are well-described by a double-exponential fit (short-dashed line). (D) Same for 50 nM octamers (*n* = 7). Panels A-D show representative curves. (E) Concentration dependence of the two time scales determined from the single-exponential and double-exponential fits for hexamers. Dashed lines are linear fits, with shaded areas representing the 95% CI of the fit. Fitted slopes are -0.60 ± 0.48 min nM^−1^ (long timescales) and -0.31 ± 0.14 min nM^−1^ (short timescales). (F) Similar for octamers. Fitted slopes are -0.28 ± 0.26 min nM^−1^ (long timescales) and -0.21 ± 0.10 min nM^−1^ (short timescales). The flow rate was 20 µL min^−1^ in these experiments.

### 4.4 Septin hexamers and octamers organize differently on membranes

The QCM-D measurements show that human septin hexamers and octamers form membrane-bound films of similar thickness (i.e., similar frequency shifts), but suggest potential differences in film structure (i.e., different dissipation shifts). To obtain direct insight in the septin film structures, we opted for atomic force microscopy (AFM), which allows high resolution imaging of membrane-bound proteins under near-native, hydrated conditions. Prior studies of septin self-assembly on membranes have often relied either on fluorescence imaging, which has limited spatial resolution, or on electron microscopy, which requires non-native conditions such as drying and metal coating (for scanning electron microscopy) or lipid monolayers (for cryo-electron microscopy) (Bertin et al. (2010); Bridges et al. (2014); Beber et al. (2019); Vogt et al. (2025); El Alaoui et al. (2025); Nakazawa et al. (2023b)). Recent studies applied AFM imaging to fly septin hexamers (Szuba et al. (2021)) and yeast septin octamers (Vial et al. (2021); Ibanes et al. (2022); Goodchild et al. (2025)). Here we apply this technique to compare human septin hexamer and octamer assemblies on membranes. Septin oligomers (100 nM) were incubated on SLBs containing 20% PS and 5% PI(4,5)P_2_. After 60 minutes of incubation, the sample was fixed with glutaraldehyde to prevent septin disruption by the AFM tip. Previous work showed that fixation does not appreciably impact septin organisation (Szuba et al. (2021)).

As shown in Figure 5A, human septin hexamers formed dense networks with patches of filaments aligned in the same direction, resembling nematic domains observed for other two-dimensional arrays of densely packed semiflexible bio-polymers (Zhang et al. (2018)). Zoomed-in images show that the hexamers form filaments that laterally associate, forming bundles that splay or join other bundles (Figure 5C). As shown in Figure 5E, a line profile across the filament bundles shows that the filaments have heights between 5 nm (as expected for single septin filaments) and 10 nm (consistent with a two-layer structure). The bundle width as observed from trough to trough in the line profile ranged from ≈ 100 nm to ≈ 500 nm (Figure 5E).

**Figure 5:**
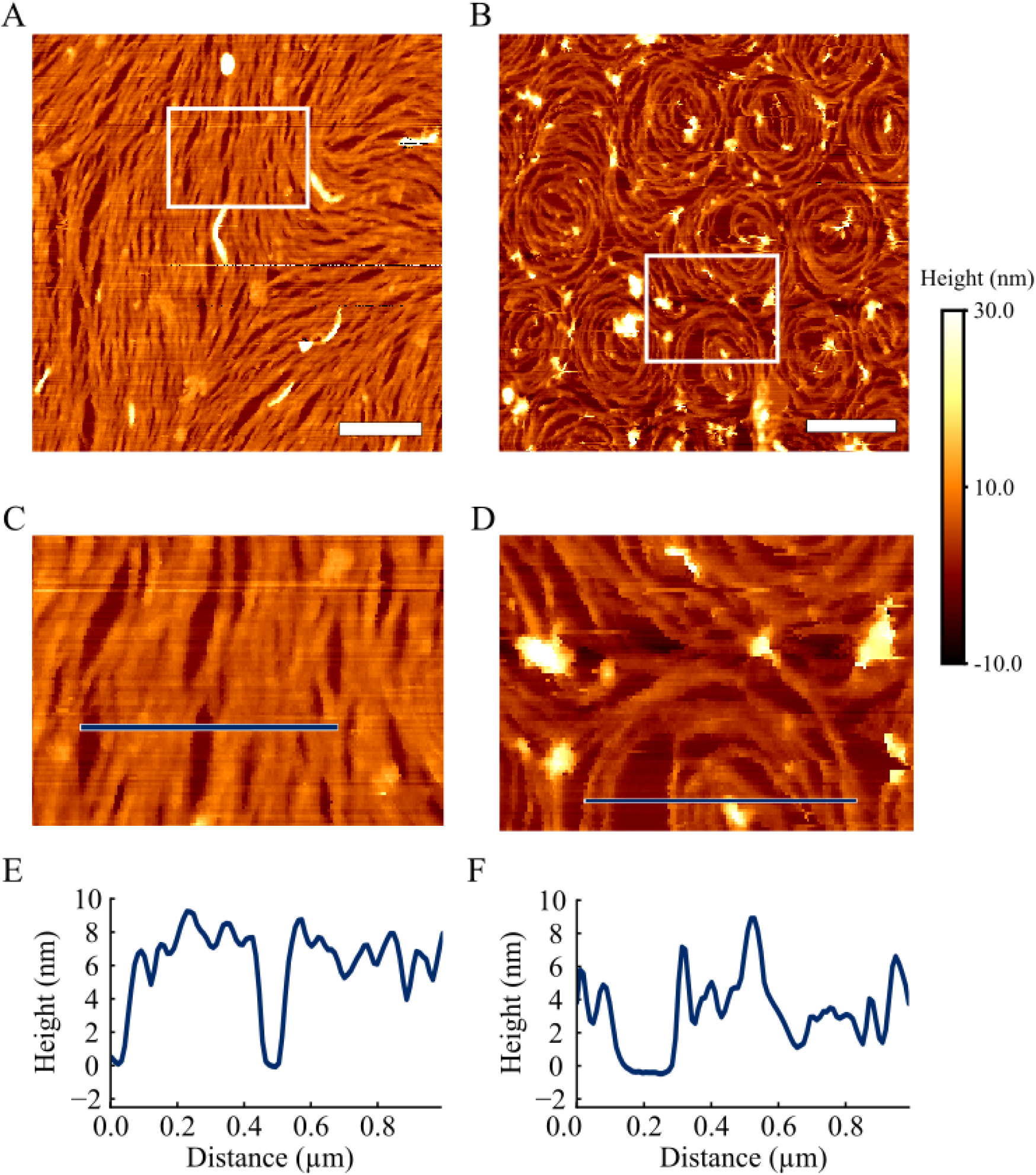
Atomic Force Microscopy (AFM) images of septin hexamer and octamer filament networks on SLB membranes. (A) Representative image of 100 nM septin hexamer filament network showing a nematic-like pattern. (B) Example image of 100 nM septin octamer filament network showing a spiral pattern. (C) Zoomed-in image of hexamers corresponding to the boxed region in panel A. (D) Zoomed-in image of octamer spirals corresponding to the boxed region in panel B. Note that the bilayer is visible in gaps in the networks. (E) Height profile along the blue line across the hexamer image in panel C. (F) Height profile along the blue line across the octamer image in panel D. Scale bars, 1 µm. Color bar represents the height, with 0 nm representing the surface of the bilayer.

When we repeated the AFM experiments with septin octamers, to our surprise we observed extended networks of vortex-like spirals in the majority of cases (Figure 5B). Across a total of 9 different experiments, we consistently observed these spiral patterns on three different experimental days (6 images). A zoomed-in image of the spirals shows that the octamer filaments/bundles are thinner than for septin hexamers (Figure 5D). In this particular example, a line profiles across one of the spirals show that the filament/bundle heights range from 4 nm to 8 nm, corresponding to maximally two filament layers (Figure 5F). Notably in one experimental session, we observed an octamer network that was more reminiscent of the nematic-like networks of hexamers (3 images) (Supplementary Figure 3A). However, the octamer filament networks showed more defects as compared to the hexamer networks, and the octamer filaments appear more highly curved than hexamer filaments. Interestingly, in one of these nematic-like network images, we observed a small cross-hatched region where one layer of filaments lays perpendicularly across the other layer (Supplementary Figure 3B). Altogether, these observations suggest that septin octamer filaments can adopt a range of curvatures and network architectures, potentially due to the incorporation of SEPT9 subunits.

A large (50 µm × 50 µm) field-of-view AFM image showed that the vortices form closely packed periodic arrays across large areas (Figure 6A). About 30 % of the vortices had an empty center (*n* = 60) (Figure 6B), while the other 70 % had a filled center (*n* = 154) (Figure 6C). This visual observation was confirmed by line profiles across spirals showing height profiles with a slight dip in height at the spiral center for the empty spirals (Figure 6D) but not the filled ones (Figure 6E). The height at the spiral outer edge (averaged over the outer 10% of the spiral diameters) was comparable for empty spirals (6.99 ± 1.31 nm) and filled spirals (6.55 ± 1.01 nm), while the center height (averaged on the inner 20% of the spiral diameter) was somewhat smaller for empty spirals (5.7 ± 1.3 nm) compared to filled spirals (7.37 ± 1.77 nm) (Figure 6G). Despite these subtle variations in filament packing within the spirals, the median diameter was comparable for empty spirals (1.02 ± 0.12 *µ*) and filled spirals (0.96 ± 0.15 *µ*) (Figure 6F). The narrow range of spiral sizes suggest a mechanism where an interplay of septin filament bending rigidity and septin-septin and septin-membrane interactions sets the size.

**Figure 6:**
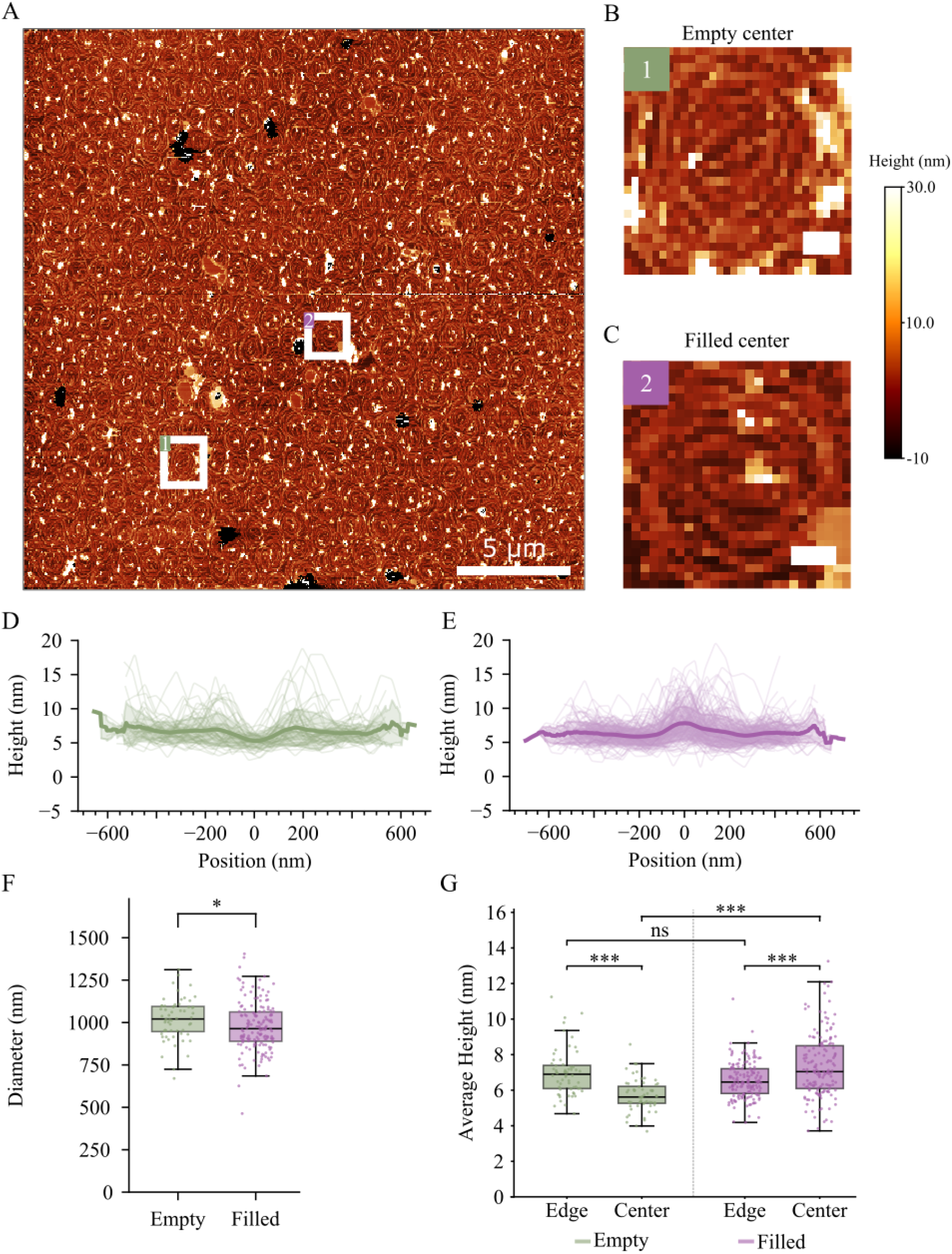
Size and height characterization of septin octamer filament spirals. (A) Large field of view AFM image of septin octamer filaments (100 nM) showing a densely packed array of regularly sized spirals. (B) Zoomed-in image of spiral with an empty interior (corresponding to area in box 1 in panel A). (C) Zoomed-in image of spiral with a filled interior (corresponding to region in box 2 in panel A). (D) Line profiles across spirals with an empty interior (*n* = 60 spirals). Light lines show individual spirals from panel A, dark line shows mean. (E) Line profiles across spirals with a filled interior (*n* = 154 spirals) from panel A, dark line shows mean. Line profiles were overlaid by shifting them so the spiral centers overlap. (F) Box plot showing spiral diameters, comparing empty (green) and filled (purple) spirals. Symbols are individual spirals, horizontal lines is the median, whiskers show standard deviation. (G) Box plot showing spiral heights at edge and center, for empty spirals (left, green) and filled spirals (right, purple). Scale bars, 5 µm for panel A and 20 nm for panel B and C. Color bar represents the height, with 0 nm representing the surface of the supported lipid bilayer. Significance is reported in p-values as *<* 0.001 (***), *<* 0.01 (**), *<* 0.05 (*) and non-significant (ns).

### 4.5 Quantitative analysis of septin network orientational order

The AFM images show that membrane-bound septin filament networks exhibit substantial orientational order. To quantify this order, we computed the filament orientations by calculating the gradients of the gray levels in the x and y directions using the OrientationPy algorithm (Rezakhaniha et al. (2012); Vasile et al. (2022)) (see schematic in Figure 7A and example workflow in **??**). Since the images are surface topographical images, the gray levels represent the height in each pixel. Hence the local orientation in each pixel gives the direction in which the height varies the least, corresponding to the direction in which the filaments in this location are aligned. From the gradient maps, we can construct vector fields representing the local orientation in each pixel by a vector, illustrated for a nematic hexamer network in Figure 7B and a spiral octamer network in Figure 7C. Using the vector maps, we can quantify the degree of alignment of the filaments in terms of the nematic order parameter, *S* (Figure 7A). This alignment metric from the liquid crystal field (Doostmohammadi and Ladoux (2022)) is commonly applied to biological systems such as actin networks and confluent epithelial cell sheets (Zemel et al. (2010); Li et al. (2025)). In a nematic liquid crystal with all filaments oriented in the same direction, *S* = 1. For an isotropic network with filaments of random orientations, *S* = 0.

**Figure 7:**
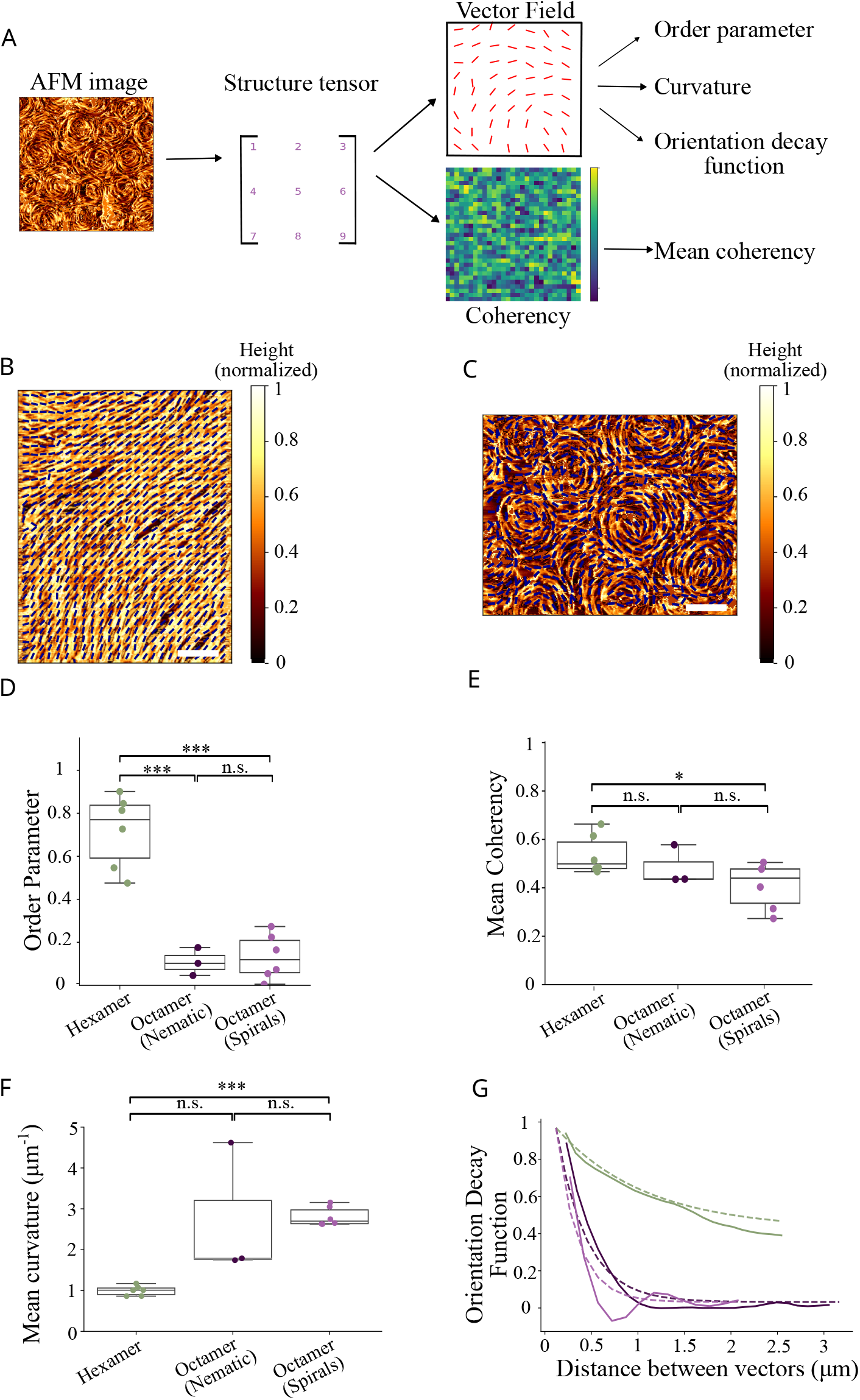
Analysis of orientational order and curvature of human septin filament networks on membranes. (A) Schematic of the analysis pipeline. Based on the AFM images, we calculate a structure tensor representing filament orientations, from which we in turn calculate vector fields and coherency. (B) Example post-processed AFM image of human septin hexamers (100 nM) overlaid with vector field showing local filament orientations within boxes with a 156 × 156 nm size. Scale bar is 1 *µm*. (C) Similar representation for octamer spirals (100 nM). Color bars represent normalized height values. (D) Box plots showing the nematic order parameter *S* of hexamers (green dots; *n* = 6), nematic octamer networks (dark purple dots, *n* = 3), and octamer spiral networks (light purple dots; *n* = 3). (E) Box plots showing the mean coherency for hexamers (green dots; *n* = 6), nematic octamers (dark purple dots, *n* = 3) and octamer spiral networks (purple dots; *n* = 6). (F) Box plots showing the mean curvatures of septin filaments (green dots; *n* = 6), nematic octamer networks (dark purple dots; *n*= 3), and octamer spiral networks (purple dots; *n* = 6). Box plots in panels D-F show average values per image (symbols) together with mean and standard deviation over all images. (G) Orientational correlation functions showing the length scale over which nematic alignment persists. Solid lines show representative examples for nematic hexamer networks (green), octamer spiral networks (light purple), and nematic octamer networks (dark purple). Dotted lines are fits to an exponential decay with offset, with average decay lengths of 0.99 ± 0.45 *µm* for hexamers, 0.34 ± 0.01 *µm* for nematic octamer networks, and 0.23 ± 0.03 *µm* for spiral octamer networks. The best-fit offset value was 0.41 ± 0.1 for hexamers. The offset was set to 0 for octamer networks. Note the oscillations in the long-distance tail of the correlation function for octamer spiral networks, indicative of regular spiral packing.

For human septin hexamer networks, we find an average order parameter of 0.72 ± 0.07 (*n* = 6), consistent with a high degree of nematic order (Figure 7D). For the human septin octamer networks, we find much smaller values for the order parameter, of 0.13 ± 0.04 (*n* = 6) for spiral networks and 0.11 ± 0.03 (*n* = 3) for nematic networks. These smaller values reflect the visual observation from the AFM images that octamer filaments are less uniaxially aligned due to filament curvature.

An alternative, more local, measure of filament alignment is the coherency, which captures the alignment of the vectors in a pixel in terms of the immediate neighbors (schematic in Figure 7A). It can take values between 0 for a disordered system to 1 for a perfectly aligned system. The mean coherency is only slightly larger for septin hexamer networks (0.54 ± 0.03, *n* = 6) than for nematic (0.48 ± 0.05, *n* = 3) and spiral (0.41 ± 0.04, *n* = 6) octamer networks, showing that hexamer and octamer filaments have similar levels of alignment on a local scale. Using the vector fields, we can also calculate the filament curvatures. For septin hexamer networks, we calculated an average filament curvature of 0.99 ± 0.25 *µm*^−1^. For octamer networks, we observed a tendency towards higher curvature (non-significant for nematic networks, significant for spiral networks), with mean values of 2.71 ± 0.95 *µm*^−1^ for nematic octamer networks and 2.81 ± 0.09 *µm*^−1^ for spiral octamer filament networks (Figure 7F). In all cases, the filament curvatures show a wide distribution within each image (Supplementary Figure 7).

To assess the spatial range over which orientational order persists across the AFM images, we calculated the orientational decay function *C*_*s*_(*r*) describing how correlations in filament alignment decrease with distance (Li et al. (2025) (schematic in Figure 7A). For septin hexamer networks, this function shows an initial exponential decay with a decay length of 0.99 ± 0.18 *µm* towards a final value of 0.5 (Figure 7F, green curves). The initial decay reflects the presence of defects in the orientational order of the filaments, known as topological defects (Doostmohammadi and Ladoux (2022)). The observation that *C*_*s*_(*r*) does not decay to zero on the size scale of the AFM images reflects the presence of long filament bundles traversing the entire image. For the octamer spiral networks, the orientational correlation function decays on a shorter length scale of 0.24 ± 0.02 *µm* and does decay to zero, indicating the absence of long-range nematic order (Figure 7F, light purple curves). Interestingly, the tail of the correlation function shows distinct oscillations, indicative of the periodic organization of the spirals. From the distances between the first and second minima in the tail, we estimate a periodicity of 1.02 ± 0.10 *µm* (*n* = 5), which corresponds to the diameter of the spirals. The orientational correlation function for the nematic octamer networks resembles that of the spiral networks, with a shorter decay length (0.33 ± 0.06 *µm*) than for hexamer filament networks and a decay to zero (Figure 7G, dark purple curves). However, unlike the spiral networks, we do not observe oscillations in the correlation function, signifying the absence of regularly sized structures. Altogether, we conclude that human septin hexamers and octamers both form organized filament networks with a high degree of local filament alignment. However, hexamer filaments are straighter than octamer filaments, giving a higher degree of long-range nematic order. These observations suggest a difference in bending rigidity and/or intrinsic curvature of filaments formed by septin hexamers versus octamers.

### 4.6 The SEPT6 and SEPT7 coiled coils are important for septin-membrane binding

The QCM-D and AFM experiments both indicate that human septins form two-layered filament networks. This is consistent with previous observations for fly septin hexamers (Szuba et al. (2021)), yeast septin octamers (Bertin et al. (2010); Jiao et al. (2020); Goodchild et al. (2025)), and human septin octamers (Nakazawa et al. (2023b)). In these prior studies, it was suggested that the first layer of septins on the membrane templates the formation of a second layer by pairing of the long C-terminal coiled coil domains on the SEPT6 and SEPT7 family septins.

Truncation of the C-termini was shown to result in the formation of septin monolayers in case of fly septin hexamers (Szuba et al. (2021)) and yeast septin octamers (Bertin et al. (2010)). To test the role of the coiled coil domains in membrane binding of human septins, we purified recombinant septin hexamers and octamers with C-terminally truncated SEPT6 and SEPT7 (referred to as ΔCC67). We first tested the membrane binding affinity of these mutants by QCM-D. To our surprise, ΔCC67-septin hexamers lost their ability to bind the membrane (Figure 8A). When we replaced the ΔCC67septin hexamers in the QCM-D flow channel with wild type hexamers, these showed normal binding in terms of frequency and dissipation shift (Figure 8A) and softness (Figure 8C), showing that the lack of binding is intrinsic to the ΔCC67-septin hexamers. AFM imaging confirmed that the ΔCC67 hexamers do not bind the membrane. By contrast, ΔCC67-octamers retained their ability to bind the membrane (Figure 8B). However, the final frequency shift after 150 minutes was reduced relative to wild-type octamers by 34%, suggesting a thinner septin film. The dissipation for the ΔCC67-octamers layer was nevertheless similar to that of WT septin octamers at equivalent septin surface coverage. Accordingly, the final softness of the ΔCC67-octamers films (0.117 × 10^−6^ Hz^−1^) was 42% higher than that of wild type octamer films (0.082 × 10^−6^ Hz^−1^) (Figure 8D).

**Figure 8:**
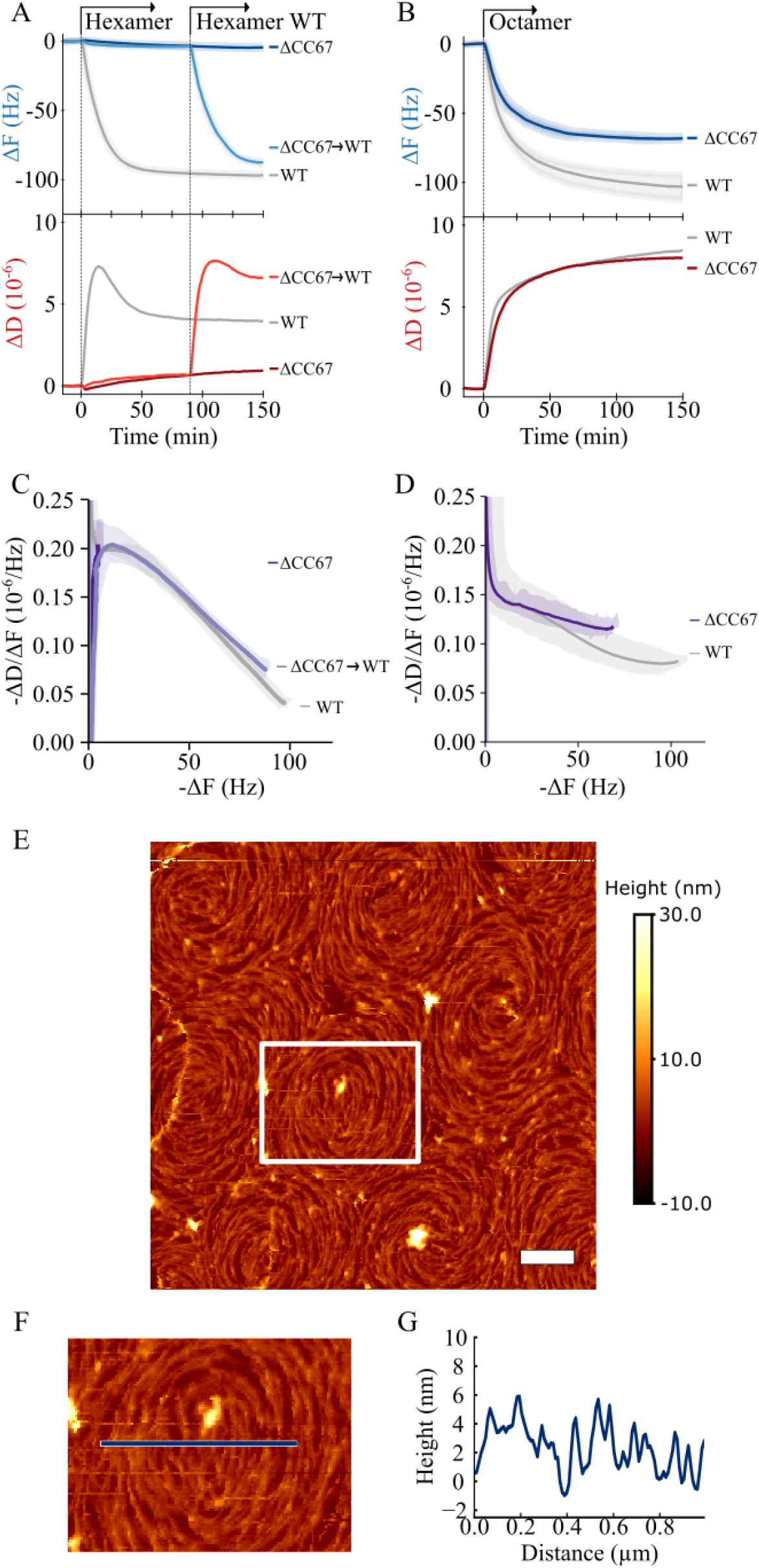
Septin coiled coils are required for membrane binding of human septin hexamers and mediate layer stacking for human septin octamers. (A) QCM-D time traces for frequency shifts (upper panel) and dissipation shifts (lower panel) obtained with a flow rate of 20 µL min^−1^ comparing 3 different experiments. Wildtype (WT) septin hexamers (50 nM, gray curves, *n* = 2) bind normally. ΔCC67-hexamers (50 nM, dark blue and red curves, *n* = 4) do not bind the SLB. WT septin hexamers flushed in to replace the ΔCC67-hexamers bind normally (light blue and red curves), showing that the lack of ΔCC67-hexamer binding is not caused by any membrane problems. (B) QCM-D data with a flow rate of 20 µL min^−1^ for human septin octamers comparing 2 different experiments. Wildtype (WT) septin octamers (50 nM, gray curves, *n* = 4) bind normally. ΔCC67-octamers (50 nM, dark blue and red curves, *n* = 4) exhibit reduced membrane binding. (C) Softness parameter ( − Δ*D/*Δ*F* ) as a function of − Δ*F*, a measure for surface coverage, computed from the data in panel A for wild type and ΔCC67 hexamers. (D) Softness parameter for wild type and ΔCC67-septin octamers calculated from the data in panel B. Shaded areas in panels C and D represent the total spread of the data. (E) Representative AFM image of ΔCC67 octamer filament network (100 nM). (F) Zoomed-in image corresponding to the boxed region in panel E. G) Height profile along the blue line across the image in panel F. Scale bars, 0.5 µm. Color bar represents the height, with 0 nm representing the surface of the supported lipid bilayer.

AFM imaging of ΔCC67-octamers on membranes revealed similar vortex-like spirals as observed for wild type octamers (Figure 8E). Moreover, zoomed-in images of spirals showed that the ΔCC67-octamer filaments within spirals had a similar organization, with a center region of straight filaments surrounded by more curved filaments (Figure 8F). Line profiles showed that the filament heights were maximally 5 nm, indicating that the ΔCC67-octamers form a monolayer – consistent with the reduced frequency shift seen by QCM-D (Figure 8G). We conclude that human septin hexamers depend on the SEPT6 and SEPT7 coiled coils for membrane binding, whereas octamers can still bind in absence of the coiled coils, likely due to the presence of the SEPT9 subunits.

## 5 Discussion

In this work, we quantitatively compared the abilities of recombinant human septin hexamers and octamers to bind and self-assemble on supported lipid bilayers (SLBs) containing PS and PI(4,5)P_2_ lipids. We showed by varying the flow rates in QCM-D measurements that membrane binding of human septin hexamers and octamers is mass-transport limited, but the steady-state septin surface coverage is flow rate-independent. This conclusion is consistent with prior QCM-D measurements for fly septin hexamers (Szuba et al. (2021)) and indicates that the intrinsic rate of membrane binding is comparatively high. The binding rate as well as the final septin coverage density increased with septin concentration. However, binding saturated even when septins were continuously perfused over the membrane, indicating that septins form a film with an intrinsically constrained thickness. The adsorption isotherms revealed positive cooperativity with effective Hill coefficients of 2.7 for hexamers and 1.2 for octamers. A similar degree of cooperativity has previously been reported for membrane binding of yeast septin octamers, where the effective Hill coefficient ranged from 1.6 in the presence of low salt (50 mM KCl, similar to our experiments) to 12.4 in presence of high salt (100 mM KCl). This positive cooperativity shows that septin-septin interactions are important for stabilising septin-membrane interactions. Consistent with this interpretation, incubation of membrane-bound septins with high-salt buffer to depolymerize septin filaments caused septin desorption from the membrane. However, we note that desorption in high-salt buffer cannot discriminate the contributions of septin-septin interaction from electrostatic effects, as both are modified simultaneously. Interestingly, septins did not desorb completely, indicating that membrane binding can stabilize filaments under high-salt conditions that would preclude polymerization in solution. Similar stabilization against depolymerization by high-salt buffer was also observed for membrane-bound yeast septin octamers (Goodchild et al. (2025)).

The maximal septin film thickness estimated from the QCM-D measurements was 17 nm for both septin hexamers and octamers, comparable to values measured previously for fly septins Szuba et al. (2021). This thickness is significantly larger than the 4-5 nm height of a single septin filament (Sirajuddin et al. (2007); Goodchild et al. (2025)), suggesting that septins form a double-layered network. AFM imaging of membrane-bound septin films showed that septin hexamers and octamers form filaments and bundles with heights of maximally 15 nm, mostly ranging between 5 and 10 nm. This observation indicates a co-existence of single filaments and stacks of two filaments layers. The film thickness from QCM-D is somewhat larger (17 nm) than the maximal heights observed by AFM. This discrepancy could be due to experimental differences between the QCM-D and AFM experiments. In QCM-D experiments, septins bind the membrane in presence of (weak) flow, whereas in AFM experiments, septins are allowed to bind the membrane from a stagnant solution. Moreover, silica-coated QCM-D sensors exhibit nanoscale roughness whereas the silicon wafers used in AFM experiments are smooth at the nm scale (Richter and Brisson (2003)). To test if flow and/or nanoscale roughness impact septin self-assembly on membranes, we developed a custom holder for the QCM-D sensor allowing us to image the QCM-D sensor after a QCMD experiment (Supplementary Figure 4). We found that septin networks on the QCM-sensors closely resembled the networks formed on silicon wafers (Supplementary Figure 5), although lateral association of septin filaments seemed slightly enhanced on the QCM-D sensor. The filament heights on the QCM-D sensor were also comparable to the heights observed on silicon wafers (Supplementary Figure 5G,H). Interestingly, we noted during the correlative QCM-D/AFM experiments that glutaraldehyde crosslinking, which is used to facilitate AFM imaging, slightly decreased the frequency shift. This could potentially signify that the fixation makes the septin films somewhat thinner, which could explain the small height discrepancy between AFM and QCM-D assays. Regardless, both assays agree that septin hexamers and octamers form double-layered filament networks. Similar conclusions were previously drawn for yeast septin octamers (Bertin et al. (2010); Beber et al. (2019); Nakazawa et al. (2023b); Goodchild et al. (2025)) and fly septin hexamers (Szuba et al. (2021)). From the AFM images, it is difficult to discern how the second layer is oriented with respect to the first layer. In electron microscopy images, a cross-hatched arrangement with 90 degree angles between filaments in the first and second layers has been reported (Bertin et al. (2010); Beber et al. (2019); Szuba et al. (2021); Nakazawa et al. (2023b)). Interestingly, in one experiment, we could observe a cross-hatched area within a swirl-like septin octamer network, with long filaments running perpendicularly across an underlying patch of filaments (Supplementary Figure 3B). In future, it will be interesting to design experiments to capture the formation process of the second layer to identify in which orientations filaments in the second layer are recruited or grow, for instance by high-speed AFM (Jiao et al. (2020); Goodchild et al. (2025)).

In prior work it was suggested that the formation of two-layered septin films on membranes relies on in-trans pairing of the long C-terminal coiled coil domains of SEPT6 and SEPT7 septins (Bertin et al. (2010); Beber et al. (2019); Szuba et al. (2021); Nakazawa et al. (2023b)). Consistent with this model, QCM-D and AFM showed that removal of the C-terminal coiled coils (ΔCC67) caused human septin octamers to form single-layer septin filament networks. This observation strongly suggests that WT octamer filaments expose their SEPT6 and SEPT7 coiled coils to the solution, facilitating the recruitment of the second layer. This interpretation is consistent with recent cryo-electron tomography of vesicle-bound human octamer septins showing two filament layers, with the first one in direct contact with the membrane and the second one spaced 12 nm apart from the first one (Nakazawa et al. (2023b)). However, to our surprise, C-terminally truncated (ΔCC67) septin hexamers lost their membrane-binding ability. One explanation could be that the coiled coils stabilize membrane binding by mediating lateral associations between septin filaments Bertin et al. (2010); Szuba et al. (2021). In solution, coiled coil-mediated filament pairing is necessary for polymerization of fly septin hexamers and yeast septin octamers Bertin et al. (2008, 2010); Szuba et al. (2021). On membranes, C-terminally truncated fly and yeast septins can polymerize, but with reduced crossbridging Bertin et al. (2010); Szuba et al. (2021). Alternatively, there may be a direct contribution of the coiled coils to membrane binding. SEPT6 coiled coils have a conserved C-terminal 18-residue region predicted to form an amphipathic helix that can insert into lipid bilayer membranes (Jiao et al. (2020); Goodchild et al. (2025)). In mammalian cells, exogenously expressed human septin hexamers lacking this helix region failed to localize to curved membrane areas and showed reduced binding to invading S. flexneri bacteria (Lobato-Márquez et al. (2021)). In case the C-terminus mediates membrane binding of septin hexamers, the filaments would likely have their coiled coils pointing downwards, which would be opposite to octamer filaments. The main distinction between octamers and hexamers is the additional presence of the SEPT9 subunits in octamers (Soroor et al. (2021); Mendonça et al. (2019)). In this work, we studied octamers containing the SEPT9i1 isoform, which has a long N-terminus with a basic region (aa 1–142) that could confer affinity for the anionic PS and PI(4,5)P_2_ lipids (Bai et al. (2013)). The SEPT9 N-terminus could drive membrane binding independently or synergistically with the coiled coils. To test these ideas, it will be interesting to compare binding and self-assembly of septin octamers harboring different SEPT9 isoforms, in particular the long isoforms SEPT9i1-i3 that share a common N-terminus versus the shorter SEPT9i4 and SEPTi5 isoforms (Kuzmić et al. (2022); Verdier-Pinard et al. (2017)). Moreover, molecular dynamics simulations could help delineate the determinants of septin-membrane binding and interplay of septin-membrane and septin-septin interactions (Lee et al. (2014); Mofidi et al. (2026)).

A very interesting outcome of our work is that human septin hexamers and octamers form differently organized networks. AFM imaging showed that hexamers form nematic-like networks that resemble organized filaments arrays seen in earlier AFM studies of fly septin hexamers (Szuba et al. (2021)) and yeast septin octamers (Goodchild et al. (2025)). By contrast, octamers predominantly formed densely packed networks of spirals with a well-defined diameter of ≈ 1 *µm*. The persistence length of human septin filaments has not yet been reported, but yeast octamer septin filaments are indeed rather flexible with persistence length values reported in a range between 3 *µm* to 12 *µm* (Bridges et al. (2014); Khan et al. (2018); Woods et al. (2021); Beber et al. (2019)). There have been previous observations of spiral-like arrangements for yeast and human septin octamers, but on curved rather than flat membranes (Beber et al. (2019); Nakazawa et al. (2023b)). The spiral shapes could derive from an intrinsically curved shape of the septin oligomers themselves and/or from interfilament contacts. There is evidence from cryo-EM imaging that septin oligomers may possess intrinsic curvature (Mendonça et al. (2021)). However, we note that the octamer filaments exhibit a range of curvatures within the spiral structures and sometimes form alternative swirl-like patterns. This suggests that the filaments are sufficiently flexible to adopt different conformations. The ability of septin octamers to form both spiral and swirl-like patterns resembles observations for filaments of FtsZ, a prokaryotic tubulin homolog (Mingorance et al. (2005); Paez et al. (2009)). In this case it was shown that a semiflexible filament model with a single set of parameters (filament curvature, persistence length, and lateral interactions) predicted the formation of both spirals and nematic textures, depending on surface coverage. Only octamers were observed to form spirals. One plausible explanation is that the SEPT9 subunits, which are absent in the hexamer, confer enhanced intrinsic curvature to the octamer by functioning as a molecular hinge (Sirajuddin et al. (2007)). Recent structural analysis suggested that the NC interface between two SEP9 septins is remarkably flexible (Castro et al. (2020)). Another possibility is that the difference stems from different interfilament interactions, originating from the distinct orientation of the septin hexamers versus octamers filaments on the membrane. To resolve these questions, additional biophysical analysis in combination with targeted mutations for human septins is needed.

Our findings reveal septin-intrinsic biophysical factors governing their binding and self-assembly on membranes. Our findings show that septins form two-layered filament networks with different network pattern formation between septin hexamers and octamers, which could explain observations of their functional roles in cellular contexts. Moreover, we showed that human septin hexamers and octamers interact differently with membranes, differing in their dependence on the SEPT6 and SEPT7 coiled coils and differing in the network structures they form. Given the complex interactome of septins it is difficult to predict how these septin-intrinsic features carry over to the complex environment of the cell. Our work provides a baseline to disentangle septin-intrinsic determinants of septin assembly from regulation by membrane properties, auxiliary cytoskeletal, and signaling proteins.

## 6 Materials and Methods

### 6.1 Plasmids and cloning

Bacterial expression plasmids for generating wild-type hexamers and octamers 9i1 were described in Iv et al. (2021) and are available through Addgene (174491, 174499, 174497, 174500). The pnEA-vH and pnCS plasmids for the bacterial expression of ΔCC67 hexamers and ΔCC67 octamers were generated using seamless cloning following the same strategy as in Ref. (Iv et al. (2021)). All primers were from Eurofins Genomics and are listed in Table 2. Primers were designed so that the encoded C-termini of SEPT6ΔCC and SEPT7ΔCC end with the sequences HYELYRRCKLEEMG and HYENYRSRKLAAVT, respectively, i.e., right after the respective *α*6 helices (underlined). All restriction enzymes were FastDigest enzymes from Thermo Fisher Scientific. All plasmids were verified by sequencing (Eurofins Genomics) after each cloning step. We have deposited all plasmids with the nonprofit repository Addgene.

**Table 1:**
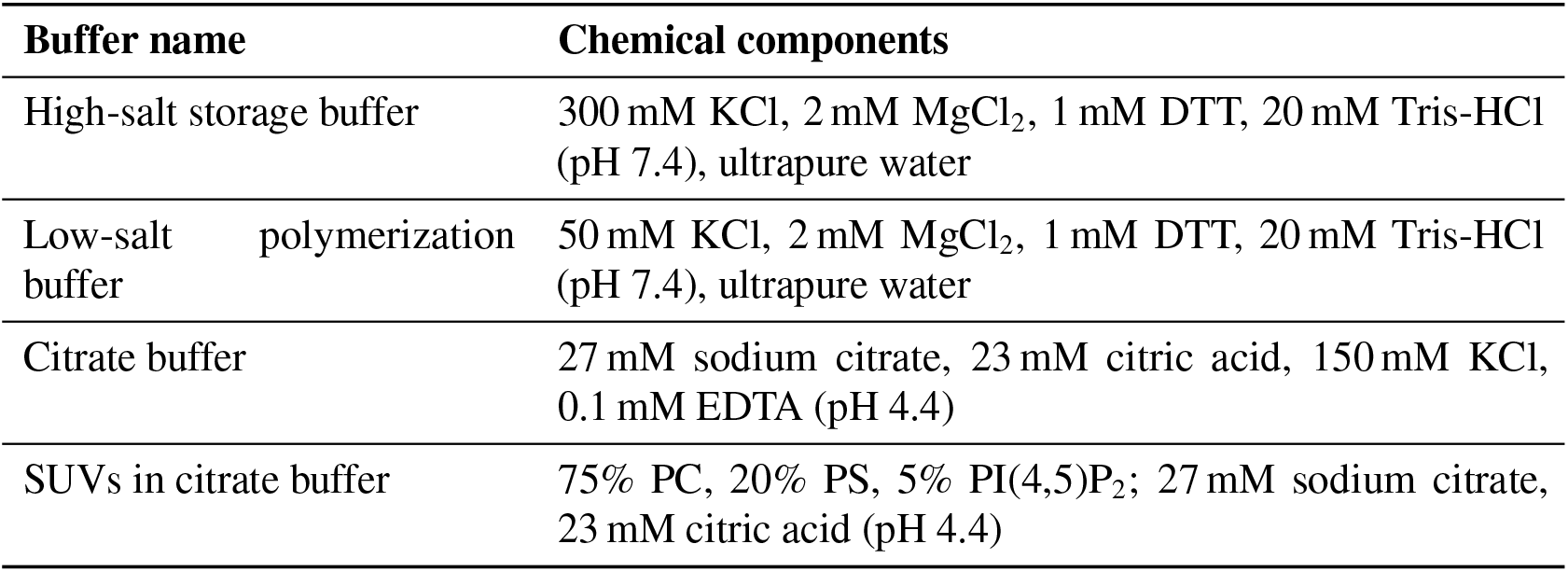
List of buffers used in septin and SLB reconstitution experiments.

**Table 2:**
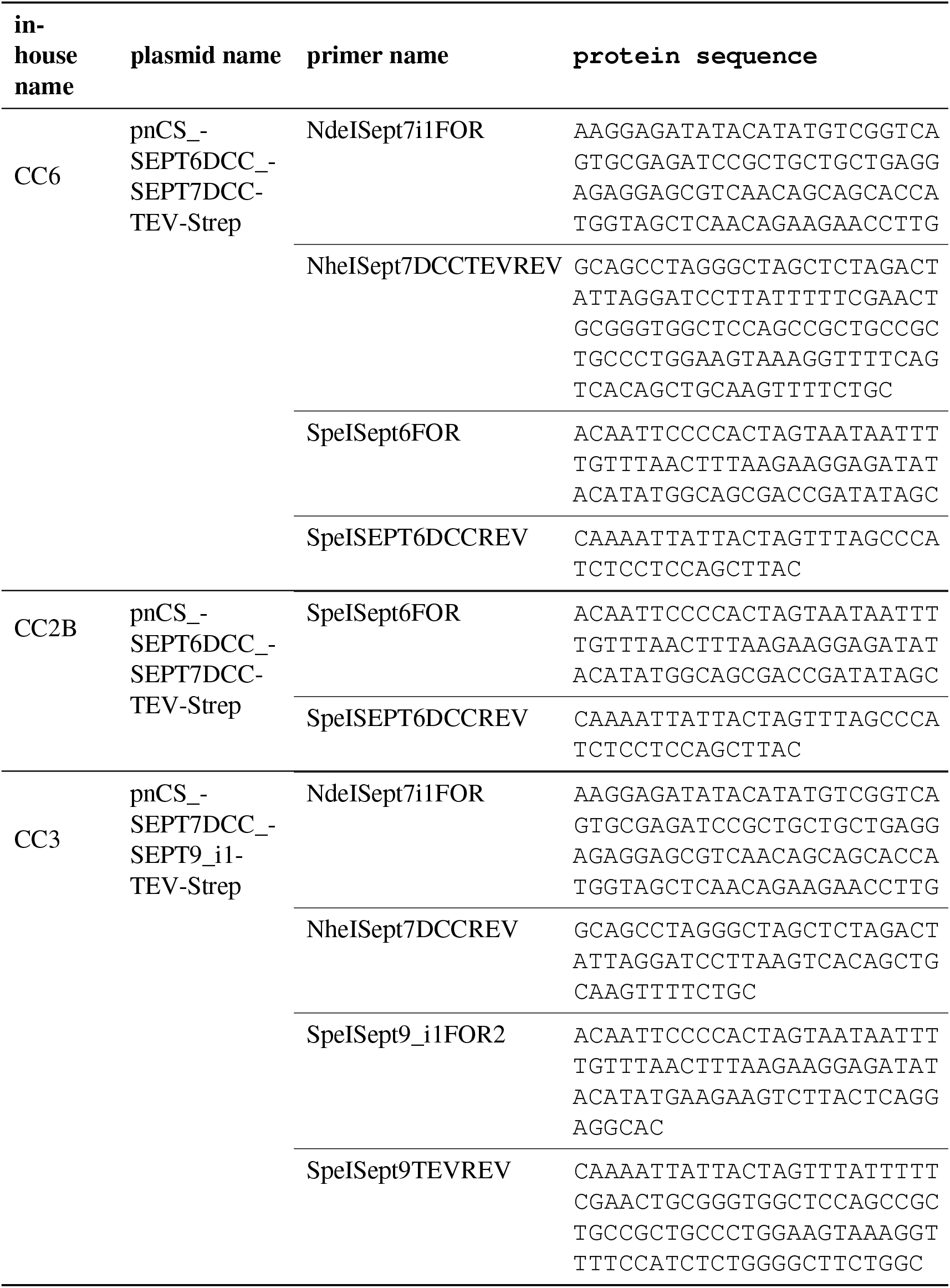
List of protein sequences for the C-truncated septin mutants.

### 6.2 Septin purification

Chemicals, unless specified otherwise, were obtained from Sigma-Aldrich. Recombinant human septin oligomers were purified in-house from BL21(DE3) *Escherichia coli* cells (New England Biolabs C2527H) using previously constructed vectors containing mono- or bicistronic constructs and a previously developed two-step affinity chromatography approach to capture septin oligomers of the correct size containing both a His6-tagged septin and a Strep-II-tagged septin (Iv et al. (2021); Castro-Linares et al. (2022)). To obtain hexamers (WT), cells were simultaneously transfected with a bicistronic pnCS vector for co-expression of full length SEPT6 and TEV-cleavable Strep-tag-II C-terminally-tagged full-length SEPT7 (Addgene nr 174499) and a mono-cistronic pnEA-vH vector for expression of full length TEV-cleavable His6 N-terminally-tagged SEPT2 (Addgene nr 174491) (Iv et al. (2021)). To obtain C-terminally truncated ΔCC67 septin hexamers, cells were simultaneously transfected with a bicistronic pnCS vector for co-expression of ΔCC SEPT6 and TEV-cleavable Strep-tag-II C-terminally-tagged ΔCC SEPT7 and a mono-cistronic pnEA-vH vector for expression of full length TEV-cleavable His6 N-terminally-tagged SEPT2 (Addgene nr 174491) (Iv et al. (2021)). To obtain octamers (WT), cells were transfected with a bicistronic pnEA-vH vector for co-expression of SEPT6 and TEV-cleavable His6 N-terminally-tagged SEPT2 (Addgene nr 174497) and another bicistronic pnCS vector for co-expression of SEPT7 and TEV-cleavable Strep-tag-II C-terminally-tagged SEPT9i1 (Addgene nr 174500) (Iv et al. (2021)). To obtain ΔCC67 septin octamers, cells were simultaneously transfected with a bicistronic pnCS vector for co-expression of ΔCC SEPT6 and TEV-cleavable His6 N-terminally-tagged SEPT2 and another bicistronic pnCS vector for co-expression of ΔCC SEPT7 and TEV-cleavable Strep-tag-II C-terminally-tagged SEPT9i1 (Addgene nr 174500).

Co-transformants were selected on LB agar plates with ampicillin (A8351-25G) and spectinomycin (BioMedicals 194540). The antibiotics were used at concentrations of 100 µg mL^−1^ and 50 µg mL^−1^, respectively, throughout the protocol. A single colony was selected to prepare an overnight preculture at 37°C with LB medium (L3022-166) containing antibiotics; the volume of the preculture was 1/50 of the final culture volume. Terrific broth (T0918-166) with antibiotics was inocculated with the preculture and incubated at 37°C. Protein production was induced with 0.5 mM isopropyl *β*-D-1-thiogalactopyranoside (IPTG, Thermo Scientific 3072587) when the optical density (OD) at 600 nm reached 2 and expression was allowed to proceed for 3 h at 37°C. Next the cells were pelleted by centrifuging the cultures at 4000 × *g* for 10 min at 4°C. Pooled cell pellets were stored at -80 °C until protein purification.

All subsequent protein purification steps were performed at 4 °C within a single day (starting from cell lysis) to minimize unnecessary exposure to proteases and contaminants. The cell pellets were resuspended in ice-cold lysis buffer (50 mM Tris-HCl, pH 8, 300 mM KCl, 5 mM MgCl_2_, 0.25 mg/ml lysozyme, 1 mM phenylmethylsulfonyl fluoride (PMSF), cOmplete protease inhibitor cocktail (1 tablet per 50 mL), 10 mg/l DNase I, and 20 mM MgSO_4_) using gentle agitation for 30 min at 4°C. Next, the cells were lysed on ice using a tip sonicator (Qsonia Q500) with 7 cycles of 30 s ON/59 s OFF pulses. The oligomers were purified by a two-step affinity chromatography procedure using an ÄKTA pure protein purification system (Cytiva). The cell lysate was first run through a His-Trap affinity chromatography column (Cytiva 29-0588-3) to capture complexes containing His6-tagged SEPT2 and the septin-containing fraction was subsequently run through a StrepTrap affinity chromatography column (Cytiva 28-9075-46) to capture complexes additionally containing StrepII-tagged SEPT7 (for hexamers) or SEPT9i1 (for octamers). The septin oligomers were finally dialyzed into a high-salt septin storage buffer (20 mM Tris-HCl, pH 7.4, 300 mM KCl, 2 mM MgCl_2_) and stored at -80°C. The high-salt buffer ensures that the septin oligomers do not polymerize.

Concentrations of septin oligomers in the stock solutions were assessed with optical absorbance measurements at a wavelength of 280 nm (Thermo Scientific, Nanodrop 2000). The absorbance measurements were converted to concentrations using extinction coefficients and molecular masses calculated from the primary amino acid sequences using ExPASy and considering two copies of each full-length septin, tags included (Iv et al. (2021); Castro-Linares et al. (2022)). The molecular masses are 291.8 kDa for hexamers and 422.5 kDa for octamers (Iv et al. (2021)). The extinction coefficients are 0.563 L g^−1^ cm^−1^ for hexamers (1 g L^−1^ = 3.4 µM) and 0.505 L g^−1^ cm^−1^ for octamers (1 g L^−1^ = 2.4 µM) (Iv et al. (2021)). For each batch, the purity and correct stoichiometry of septin subunits were checked using denaturing electrophoresis (SDS-PAGE) using 4–20% precast polyacrylamide gels (Mini-PROTEAN TGX Gels from Bio-Rad, 4561095) stained with InstantBlue Coomassie stain (Expedeon, ISB1L). Molecular mass markers were Precision Plus Protein All Blue Standards from Bio-Rad (1610373). For both hexamers and octamers, clear bands were visible for all the individual septin proteins (Supplementary Figure 8A and B) and for the ΔCC67 septin hexamers and octamers (Supplementary Figure 9A and B). In addition, the integrity and stability of the complexes (at a concentration of 12.5 nM in storage buffer) for each batch were examined by mass photometry (OneMP, Refeyn) (Castro-Linares et al. (2022)). The hexamer preparations contained a majority fraction (88%) of hexameric complexes (Supplementary Figure 8C) while the octamer preparations contained 33% of octameric complexes (Supplementary Figure 8D). The remaining peaks indicate partially formed complexes (dimers and tetramers).

### 6.3 Preparation of septin polymerization pre-mix solutions

On the day of the experiments, septin oligomer aliquots were thawed and kept on ice. First, the septin oligomer stock was diluted in high-salt septin storage buffer to obtain a high-salt pre-mix solution at a septin oligomer concentration 6-fold higher than the final target concentration (Iv et al. (2021)). Next, an ≈ 5-fold concentrated polymerization buffer without KCl (58.3 mM Tris-HCl pH 7.4, 5.8 mM MgCl_2_, 5 mM DTT and 5 mM GTP) was made. Finally, the septin high-salt pre-mix solution was mixed with this buffer solution to generate septin solutions in low-salt septin polymerization buffer conditions (20 mM Tris-HCl pH 7.4, 50 mM KCl, 2 mM MgCl_2_, 1 mM DTT, 1 mM GTP) at final septin oligomer concentrations of 200 nM (for AFM, later diluted to 100 nM in low-salt buffer) or 6.25, 12.5, 25, 50 or 100 nM (for QCM-D). This solution was immediately added into the AFM well or perfused into the QCM-D system.

### 6.4 Preparation of small unilamellar vesicles (SUVs)

Lipids were purchased from Avanti Research. PC (2-di-(9Z-octadecenoyl)-*sn*-glycero-3-phosphocholine, Merck 850375C) and PS (1,2-di-(9Z-octadecenoyl)-sn-glycero-3-phospho-L-serine (sodium salt), Merck 840035C) were received in chloroform and stored in glass vials under argon. PI(4,5)P_2_ (L-*α*-Phosphatidyl-D-myo-inositol-4,5-bisphosphate, Triammonium Salt, Porcine Brain PI(4,5)P_2_, Merck 840046P) was suspended in a 20:9:1 v/v mixture of chloroform:methanol:ultrapure water and aliquotted for storage in glass vials under argon. The lipids were diluted in the same 20:9:1 v/v solvent mixture in a fixed molar ratio of 75:20:5 of PC:PS:PI(4,5)P_2_ and then dried under a stream of nitrogen gas. The lipid film was placed in a vacuum desiccator for 2 h to remove any residual solvent. The lipids were then resuspended at a total concentration of 0.25 mM in an acidic citrate buffer (27 mM sodium citrate, 23 mM citric acid, 150 mM KCl, 0.1 mM EDTA, pH 4.4). The acidic pH promotes the formation of homogeneous and fluid supported lipid bilayers by reducing the net charge on the head group of PI(4,5)P_2_ from − 4 to − 3 (Braunger et al. (2013)). The lipid solution was vortexed using 4 cycles of 30 s vortexing/5 min waiting and subsequently sonicated in a bath sonicator (Sonopuls HD 2070.2 with a BR 30 cup, Bandelin) for 60 min at 10% amplitude with 5 s on/off pulses. The resulting solution of small unilamellar vesicles (SUVs) was wrapped with parafilm and stored at 4 °C for up to 7 days as in previous studies (Braunger et al. (2013); Szuba et al. (2021); Iv et al. (2021)). For QCM-D, the SUVs were diluted 4-fold into citrate buffer immediately before use. For AFM, the SUVs were directly added to the silicon wafer.

### 6.5 Quartz crystal microbalance with dissipation (QCM-D)

Before each QCM-D experiment, silicon dioxide coated QCM-D sensors (QSX303, Biolin Scientific) were cleaned by submersion for 30 min in a solution of 2% sodium dodecyl sulfate (SDS) in MilliQ (MQ) water. The QCM-D sensors were rinsed with ultrapure water, dried with a stream of nitrogen gas, and treated by UV/ozone (Ossila, L2002A3-EU) for 30 min to render the surface hydrophilic. The cleaned QCM-D sensors were immediately placed in the flow chambers of the QSense E4 QCM-D system (Biolin Scientific). The outlet tubing was connected to a syringe pump (AL-1600, World Precision Instruments), to allow a controlled flow of liquid over the sensor at a flow rate of 20 µL min^−1^ unless otherwise stated. All experiments were conducted at 22 °C. First, the low-salt polymerization buffer was perfused into the QCM-D channel for 10 min to determine the baseline values of the frequency shifts Δ*F*_*i*_ for the fundamental (*i* = 1 with resonance frequency of 5 MHz) and harmonics (*i* = 3, 5, 7, 11 and 13, corresponding to resonance frequencies of approximately 15, 25, 35, 45, 55 and 65 MHz) and corresponding dissipation values Δ*D*_*i*_. Then, the system was perfused with citrate buffer (10 min) before adding the SUVs in citrate buffer. Once the SLB had formed (as monitored from the Δ*F*_*i*_ and Δ*D*_*i*_ signals (Richter et al. (2003); Baumann et al. (2010)), the system was perfused with citrate buffer (5 min), then low-salt polymerization buffer (10 min), and septin polymerization pre-mix solution (maximally 150 min). Finally we tested the reversibility of septins in low-salt polymerization buffer (10 min) or high-salt septin storage buffer (20 min) and tested for perturbations of the SLB by protein denaturation with 6 M guanidinium chloride (10 min) followed by washing with low-salt polymerization buffer (20 min).

### 6.6 QCM-D data processing and analysis

The reported frequency shifts were normalized with the harmonic number (Δ*F* = Δ*f*_*i*_*/i*), since for adsorption of thin and rigid layers, the true frequency shift is proportional to the harmonic (Sauerbrey (1959)). In this work, only the measurements for the fifth harmonic are shown, as this proved to be the most stable overtone. A Gaussian smoothing kernel with standard deviation of 10 data points ( ≈ 15 s) was used to remove high-frequency noise in the frequency and dissipation signals.

We use the Sauerbrey equation to relate the frequency shift (Δ*F* ) to the bound septin mass:

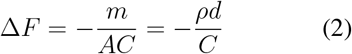

where *m/A* is the areal mass density, *d* the layer thickness, *ρ* the layer mass density, and *C* the mass sensitivity constant of the sensor. To calculate septin layer thickness, we assumed that the volume fraction *ϕ* of septins in the septin layer is 10%. While this number may seem low, even in protein crystals, the volume fraction of solvent often exceeds 50% (Kantardjieff and Rupp (2003)). Given a protein density *ρ*_*p*_ of 1.4 g cm^−3^ (Fischer et al. (2004)) and solvent density *ρ*_*s*_ of 1.0 g cm^−3^, the mean film density is

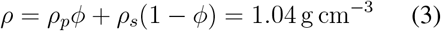

For calculating the septin layer thickness (*d*), we use the Sauerbrey equation (see Equation 2):

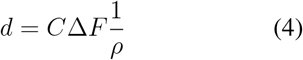

The uncertainty *σ*_*d*_ in the thickness estimate was computed by propagating the measurement error (*σ*_*C*Δ*F*_ ) and the potential error in our assumption of protein film density (*σ*_*ρ*_):

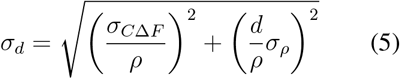

As the septin volume fraction in the film is likely between 0% and 20%, we take *σ*_*ρ*_ as the standard deviation of a uniform distribution between densities of 1.0 g cm^−3^ and 1.08 g cm^−3^.

To quantify the dependence of septin adsorption on bulk septin concentration *S*, we performed a least-squares fit of a Hill function to the the final near-steady-state absorption levels Δ*F*_ss_:

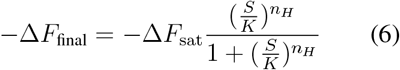

with the frequency shift at saturation *F*_sat_, effective dissociation constant *K*, and Hill coefficient *n*_*H*_ as fit parameters.

To analyze the kinetics of septin binding, we performed two types of phenomenological analyses. First, we estimated the septin binding rates from the time derivatives of the frequency shift data for each individual experiment. The dependence of the rate of change of the frequency shift on septin concentration (Figure 3C and D) suggests that, to a first approximation, the septin binding rate is a linear function of septin concentration *S*. Hence the change in number of septin oligomers *N* per unit area *A* (in the initial phase of binding where unbinding is negligible) is given by:

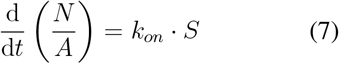

Previous work reports that in sparse monolayers of globular proteins and spherical virions, the hydrodynamically trapped solvent can contribute between 60% and 90% of the total monolayer mass (Reviakine et al. (2011)). This value is not known for sparse septin monolayers in the initial stages of binding, but to a first approximation, we take the septin and trapped solvent mass fractions to be 10% and 90%, respectively, as any exposed coiled-coils likely trap a relatively large amount of solvent. With septin mass fraction *ω*_*s*_ = 0.1, the septin number density *N/A* can be related to the total mass density *m/A*:

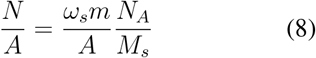

with *M*_*s*_ representing the molecular weight of a septin oligomer and *N*_*A*_ Avogadro’s number constant. Using the Sauerbrey equation (Equation 2), we can relate the mass density *m/A* to the frequency shift Δ*F* :

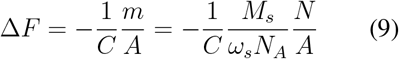

Combining Equations 9 and 7, the on-rate *k*_*on*_ becomes:

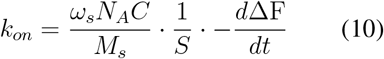

where 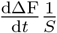 is obtained as the slope of the linear fits in Figure 3C and D. Note that this binding rate estimation strongly depends on the actual mass fraction of septins in the septin layer. Furthermore, our approach underestimates the true association rate, as we find that the binding of septins is mass-transport limited. Instead, it represents the effective association rate for a given condition.

The second method to determine the kinetics of septin-membrane binding consisted of fitting the QCM-D time traces with a sum of up to three exponentials (with *n* ∈ 1, 2, 3):

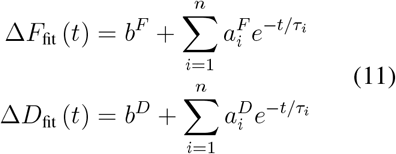

We simultaneously fitted the frequency and dissipation shift data, such that the timescales *τ*_*i*_ are shared. Note that the amplitudes 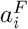 and 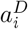 and offsets *b*^*F*^ and *b*^*D*^ (which we did not interpret) are specific to either the frequency or dissipation shift, respectively. To identify optimal fit parameters, we performed least-squares regression with residuals:

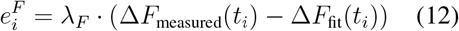

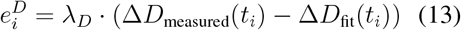

Where 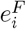 and 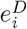 are the residual at time point *t*_*i*_ for Δ*F* and Δ*D*. Using least square regression, we minimize

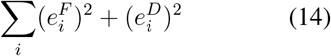

The weights *λ*_*F*_ (in units of Hz^−1^) and *λ*_*D*_ (in units of 1*/*10^6^) determine the relative contributions of the frequency and dissipation curve to the residual. To weight the frequency and dissipation residuals equally, we set *λ*_*F*_ = 1 and *λ*_*D*_ = 20, as the magnitude of − Δ*F* (in Hz) is typically 20 times larger than the magnitude of − Δ*D* (in units of 10^−6^). We compared the mean squared residuals from single-, double- and triple-exponential fits, which showed that single-exponential curve fits captured the time traces at low septin concentrations while double-exponential curve fits were needed at higher concentrations of hexamers (Supplementary Figure 2A) and octamers (Supplementary Figure 2B).

### 6.7 Atomic force Microscopy (AFM)

Ultraflat silicon wafers (Silicon Prime wafer, N/arsenic) were segmented with an engraving pen (Sigma) into pieces of 1 × 1 cm^2^ for each new AFM experiment. The wafers were rinsed with ethanol and ultrapure water, dried using a stream of nitrogen gas, and rendered hydrophilic by UV/ozone (Ossila, L2002A3EU) treatment for 30 min. The cleaned silicon wafers were immediately glued to a microscope slide (Epredia) with two-component glue (Bison kombi snel, EAN 8710439014142). After forming a 1 cm diameter boundary ring of grease (Sigma Aldrich, Z273554), we deposited 100 µL SUVs in the well. After 30 min of incubation at room temperature, excess SUVs were removed by washing 6 times with acidic citrate buffer, each time removing 50 µL of the solution on the wafer and replacing it with 50 µL fresh solution. The buffer was then exchanged for the low-salt polymerization buffer using 6 sequential removals and replacements of 50 µL. Finally, 100 µL of the low-salt buffer was replaced with the septin polymerization pre-mix solution (containing 200 nM septin oligomers), resulting in a final septin concentration of 100 nM. After 1 hour of incubation at room temperature, unless otherwise stated, the sample networks were chemically cross-linked with 1 wt% glutaraldehyde (GTA) in low-salt polymerization buffer. Crosslinking was previously shown to prevent disruption of septin filaments on membranes during AFM-scanning without significantly altering the protein structures (Szuba et al. (2021)). Finally, the sample was washed with low-salt polymerization buffer before imaging. Samples were imaged with a JPK-Bruker Nanowizard 4XP using a V-shaped ScanaSyst Fluid+ cantilever with a nominal spring constant of 0.7 N/m and tip radius of 2 nm. The images were acquired in quantitative imaging mode with a setpoint force of 1.5 nN, a z-length of 50-100 nm, and a scan rate of 5 ms per pixel for 512 × 512 pixels. We recorded square regions of interest ranging in size from 5 × 5 µm up to 50 × 50 µm.

Before any analysis, AFM images were post-processed in Gwyddion (Nečas and Klapetek (2011)). First, large imaging artifacts were cropped. Any large structures that were judged to be aggregates or debris were manually masked. Second, the scan rows were aligned with a second-degree polynomial fit to correct the drift to a uniform common baseline. The height (z-position) of the top surface of the lipid bilayer membrane was determined by masking the filaments based on height and then inverting the masked-filament height. Then the masked background was leveled to a mean height of zero, thus setting zero height equal to the surface of the membrane visible in gaps between septin filaments.

The orientational order of the septin filament networks was determined with the open source OrientationPy package (Vasile et al. (2022)). First, the text files containing the post-processed AFM images were further corrected for scan line artifacts in Python. This was done in the 2D Fourier domain, where vertical frequency lines corresponding to scan line artifacts were subtracted. The bandwidth of these lines was set to 6 pixels^−1^, which was the value at which the line errors got reduced, while no additional artifacts were introduced to the image. Next, the height values were normalized based on the minimum and maximum heights for each image to ensure consistency across images. We divided the 512 × 512 pixels images (5 µm × 5 µm in physical dimensions) into square regions of 16 × 16 pixels and for each region we computed the structure tensor, with components *J*_*xx*_, *J*_*xy*_, *J*_*yy*_, using OrientationPy. The structure tensor determines the general orientation of filaments in the 16 × 16 pixels region based on the direction in which there is minimal variation in height. The preferred direction was converted into an angle *θ*, where an angle of 0°represents the horizontal direction. To calculate the vector describing each 16 × 16 pixels region within an image, we chose its starting point as the center of the square region, and its direction along the angle extracted from the structure tensor. The length of the vector was chosen as the coherency value within the box.

We calculated two distinct measures of the degree of orientational order in each image. The first measure is the coherency calculated from the eigenvalues of the structure tensor, representing the maximum and minimum variance of the orientation (Vasile et al. (2022)). The coherency is a scalar measure that quantifies the “sharpness” or “directional strength” of the dominant orientation within each image window (16 × 16 pixel regions in our analysis). It ranges from 0 for randomly ordered structure to 1 for perfect alignment along a single dominant direction. The second measure is the nematic order parameter *S*, used widely to describe the global degree of uniaxial alignment of nematic liquid crystals (Doostmohammadi and Ladoux (2022)). We calculate *S* from the angles extracted from the structure tensors as follows:

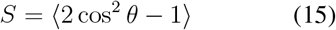

In a perfectly ordered nematic liquid, in which all filaments are oriented in the same direction, *S* = 1, while *S* = 0 for an isotropic network in which the filaments have random orientations. To assess the range over which the nematic alignment persists, we computed an orientational correlation function *C*_*s*_(*r*), which describes how the dominant orientation at one point in the image compares with the orientations at points located at a radial distance *r* (Li et al. (2025)). For efficiency, we used FFT auto-correlation in Python, which computes the correlations for all distances at once, using spatial frequencies instead of going over each vector pair individually. The orientational correlation function for each image was fitted to the sum of a single-exponential exponential decay with decay length *ξ* plus an offset *C*_∞_:

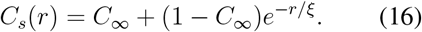

For octamer spiral and nematic patterns, *C*_∞_ was set to zero, while for hexamers, *C*_∞_ was a fit parameter. For the fit values for decay lengths (and offsets for hexamers) per image, we computed mean values and 95% confidence interval.

From the vector field, we also computed the curvature from the changes in vector orientations between the central vector in each 16 × 16 pixels regions and the vectors in the 8 adjoining regions. For the horizontal and vertical neighbors, the curvature was calculated using the formula:

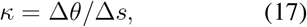

where Δ*θ* represents the angle difference between the central vector and the neighbor vectors, and Δ*s* represents the physical spacing.

For diagonal neighbors, the same formula was used, but with a scaling factor of 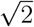. The curvature of the central vector was finally determined by the average of the resulting 8 curvature values. To compare different conditions, we computed the mean curvature per image.

In Gwyddion, the largest image (field of view of 40 µm by 40 µm, containing *n* = 258 spirals) was used to estimate the size and height distribution of spirals. For each spiral, a line height profile was measured across its shortest axis (hereafter defined as x-axis). First, the spiral diameters were estimated using a custom script in Python from the total length of the line profiles. Second, the differences in heights between spiral periphery and center were investigated. To this end, spirals were first classified into spirals with an empty or filled center by centering the line profiles at *x* = 0, normalizing the height values by the maximum height, and classifying spirals as empty (*n* = 60) if the mean normalized height of the 10% central region was below a 35% threshold or filled (*n* = 154) otherwise. Spirals with a central region having a linear regression slope of center 10% of the normalized height vs *x* greater than 0.40 (dimensionless, normalized by the center band width) were removed from the analysis, since the steep slope indicated the profile could not be at a trough or peak. After classification, we determined absolute height values (nm) of the central portion of the spirals (central 20% of each line profile) and edge region (the outer 10% from either end). Welch’s t-test was performed to compare the spiral diameters of the two populations. An ANOVA oneway was used to compare the heights of the spiral regions, the p-values were determined with the Scheffe post-hoc method.

### 6.8 Correlative AFM-QCM-D experiments

To allow AFM imaging of QCM-D sensors after QCM-D experiments, we designed a custom QCM-D sensor holder to keep the networks hydrated after removal from the closed QCM-D channels. The 3D holder consisted of two parts that were fabricated using the Fusion360 3D-CAD software (3D object geometry file available upon request). Both parts were 3D-printed from polyethylene terephthalate glycol (PETG) by fused deposition modeling with the Original Prusa XL or Original Prusa MK4 3D printer using a 0.4 mm nozzle. Upon assembly, a central chamber is formed in which the QCM-D sensor is held between two O-rings (QSense standard o-ring QSS001) (see Supplementary Figure 4). A hole in the top cover of the sensor holder allows access to the binding surface of the sensor. During AFM measurements, the two halves are held together by slide holder clips pressing down on the two flat extensions, creating a watertight seal.

We first performed QCM-D experiments, monitoring septin assembly on SLB-coated QCM-D sensors as described above. Once assembly reached steady state (100-150 min after septin perfusion), we perfused the channel with glutaraldehyde (0.1% in low-salt polymerization buffer) for 1 min to cross-link the septin network followed by a 5 min wash with low-salt buffer. The flow was turned off and the QCM-D sensor was transferred to the custom designed sensor holder for AFM imaging.

## Acknowledgments

We thank Jeffrey den Haan and Sonam Marapin for protein purification and Allard Katan for help with AFM experiments and data analysis. We thank Abigail Roberts, Erin Tait, James Hooper, Jennifer Barber, and Itzel Garcia Monge from the Richter lab for training and scientific discussions. We furthermore thank Aurelie Bertin, Koyomi Nakazawa, Stephanie Mangenot, Gerard Castro Linares, Timon Idema and Felix Frey for helpful discussions. This work was supported by startup funds from the Bionanoscience Department and the TU Delft Excellence Fund (GK), Research Grants BB/X007278/1 and BB/X00158X/1 from UK Research and Innovation (BBSRC) (R.P.R.) and the Centre national de la recherche scientifique (CNRS; MM).

## Author Contributions

**S.R**., **G.H.K, M.M. and R.P.R**. conceived and designed experiments; **M.M and S.O.M** designed and cloned plasmids; **S.R**., **W.d.R, and A.v.H** performed experiments; **S.R**., **W.d.R, A.G.M. and R.T**. developed computational analysis tools; **S.R**., **W.d.R, A.G.M, R.T, and R.P.R** analyzed data; **S.R**., **W.d.R, A.G.M, and R.T** prepared the digital images; **S.R**., **W.d.R, A.G.M, and G.H.K**. wrote the manuscript with input from all authors.

## Data Availability

All data supporting the findings of this study are available within the paper and its supplementary information files. Raw data is available from the corresponding author upon reasonable request.

## 9 Supplementary Figures

