## Supplementary figures for "Quantitative biophysical analysis of human septin hexamer and octamer self-assembly on model membranes"

A

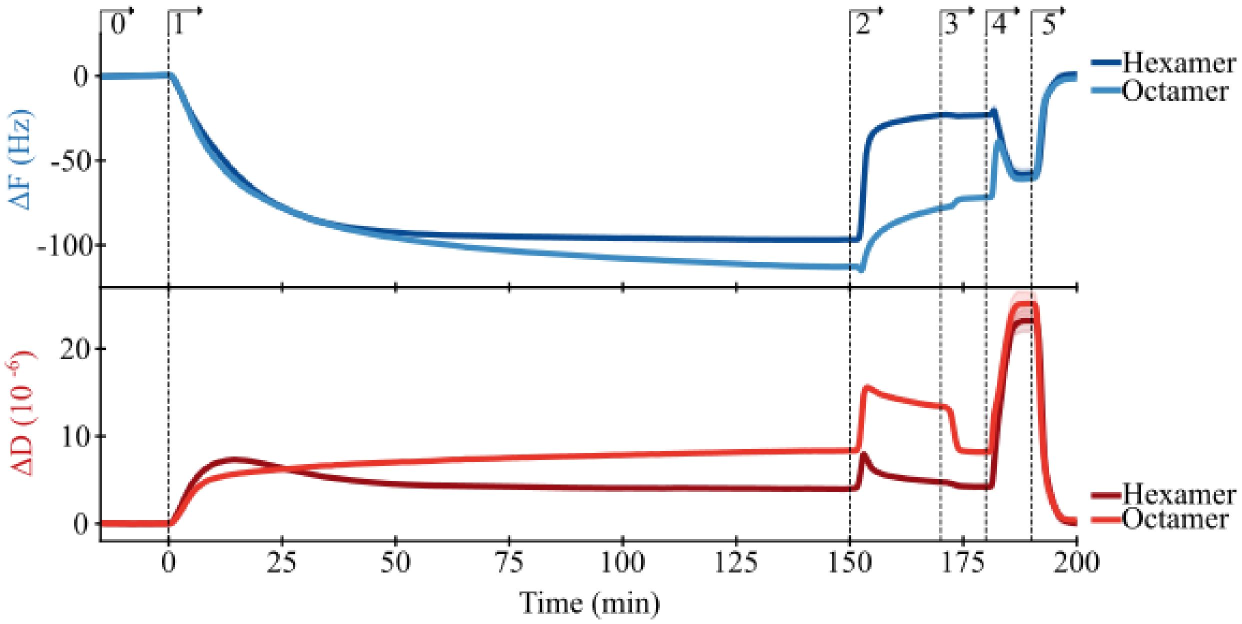


**Supplementary Figure 1:** Control QCM-D experiments showing that septins largely desorb from the membrane in high salt buffer and septin binding does not disrupt the SLB. (A) Frequency shifts (top) and dissipation shifts (bottom) for human septin hexamers ($n$ = 2) and octamers ($n$ = 2). Numbered arrows show sequential perfusion with: (0) low-salt polymerization buffer; (1) 50 nM septin oligomers; (2) high-salt buffer to depolymerize and desorb septin filaments; (3) low-salt polymerization buffer to compare to buffer baseline; (4) 6 M guanidinium chloride to denature remaining protein; and (5) low-salt polymerization buffer to return to the SLB baseline. The changes in $\Delta F$ and $\Delta D$ accompanying the transitions from step 4 to 5 and step 3 to 4 are largely due to changes in the viscosity and/or density of the solutions used rather than surface effects. Note that there is a transient increase in dissipation on rinsing with high-salt buffer (step 2) for the octamers but not the hexamers. This suggests that the membrane-bound octamer film undergoes some reversible salt-dependent changes in structure (and thus viscoelastic properties). Lines represent the mean and shaded areas the data spread. For buffer compositions: see Table 1.

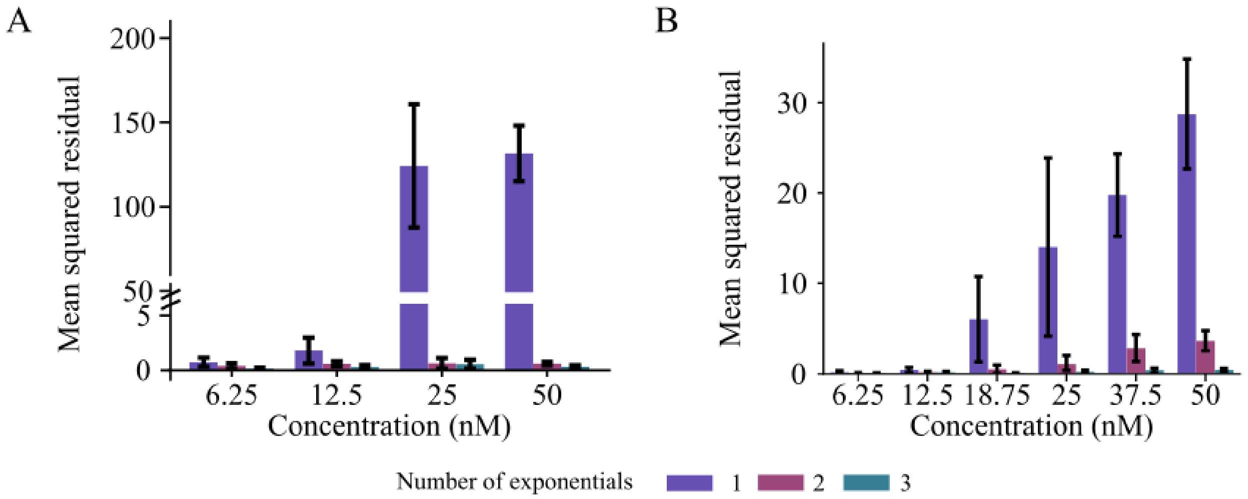


**Supplementary Figure 2:** Average mean squared residuals of the exponential fits to extract time scales of human septin binding from QCM-D data. (A) Mean squared residuals of single-, double-, and triple-exponential fits for hexamer binding curves (50 nM). (B) Mean squared residuals of single-, double-, and triple-exponential fits for octamer binding curves (50 nM). Error bars are standard deviation. Note that the QCM-D data are the same data as in Figure 3A and B, obtained at a flow rate of 20 μL $min^{-1}$.

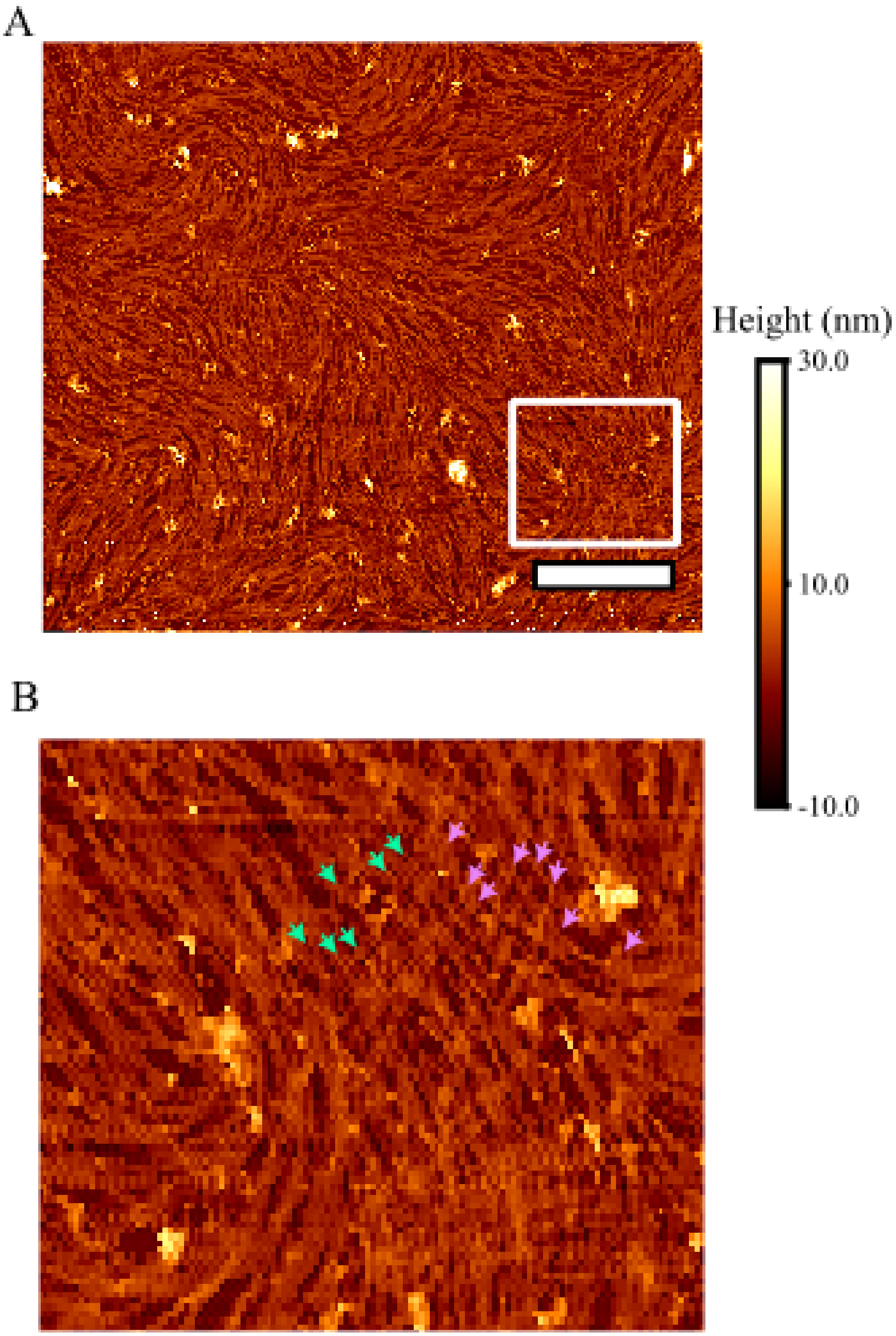


**Supplementary Figure 3:** Alternative nematic network morphology observed for human septin octamers (100 nm) on SLBs. (A) A representative AFM image of a filamentous septin octamer network forming a nematic pattern. This pattern somewhat resembles the nematic septin hexamer networks, but with more defects and more highly curved filaments. (B) Zoomed-in image of the boxed region from panel A, showing a cross-hatched pattern of filaments as indicated by the arrows showing one set of filaments (cyan arrows) crossing another set of filaments (magenta arrows). Scale bars, 1 µm. Color bar represents the height, with 0 nm representing the bilayer surface.

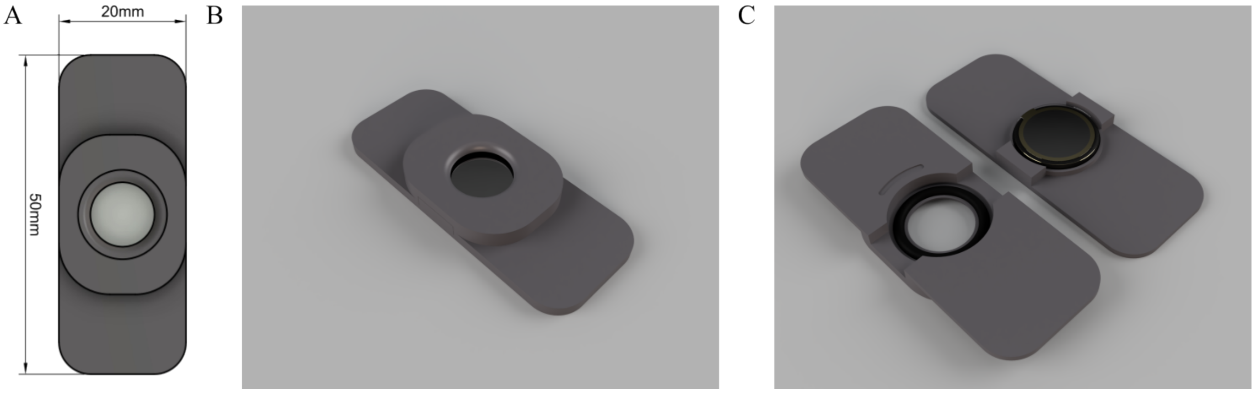


**Supplementary Figure 4:** Design of the custom 3D-printed sample holder for correlative AFM/QCM-D experiments. (A) Top-down view of the sensor holder with dimensions as shown. (B) Assembled view of the sensor holder. (C) Disassembled view of the sensor holder. Note the sensor lying in between the two rubber O-rings, which are slotted in circular depressions.

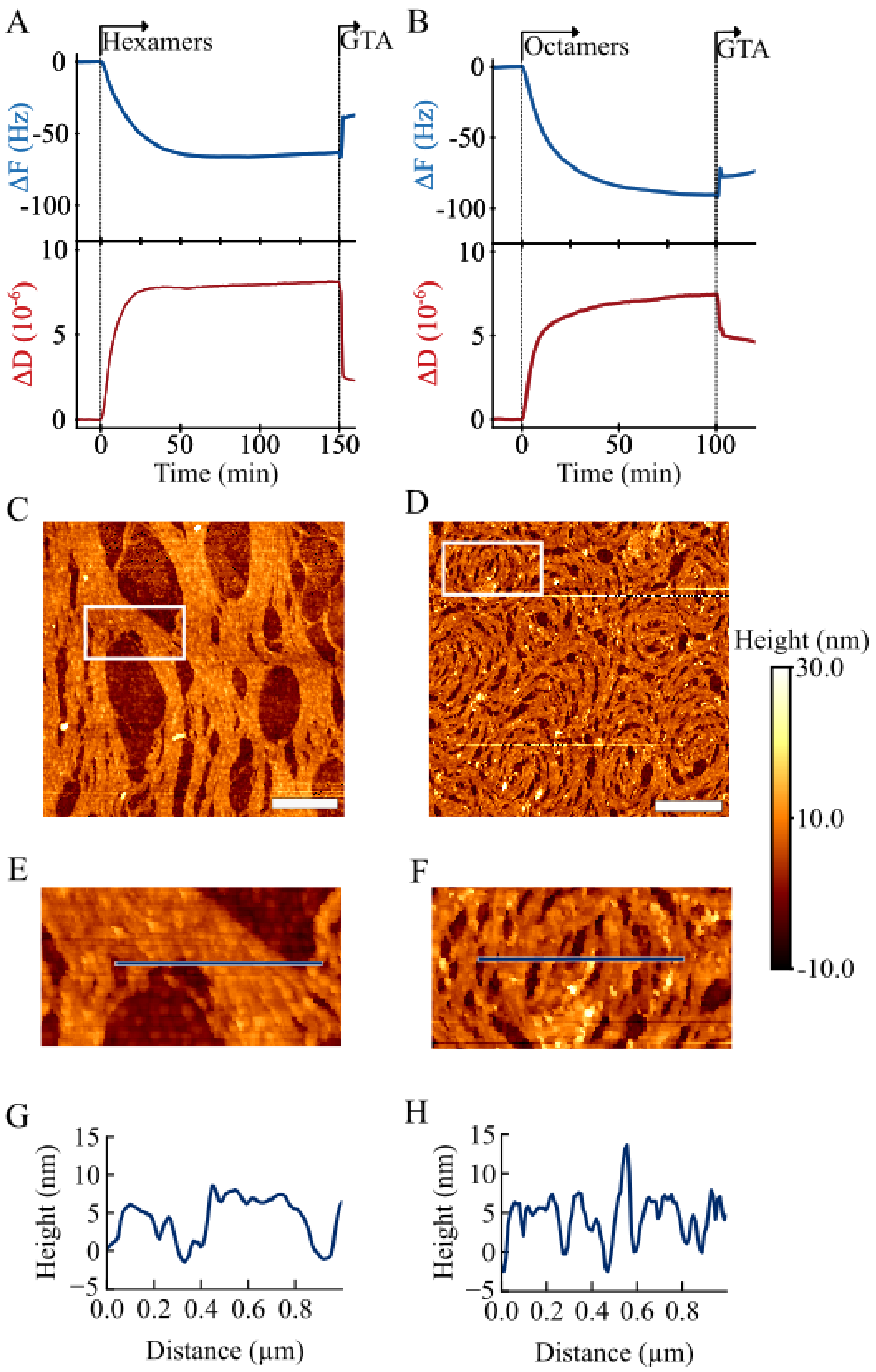
A
Hexamers
GTA
ΔF (Hz)
ΔD (10⁻⁶)
Time (min)
B
Octamers
GTA
ΔF (Hz)
ΔD (10⁻⁶)
Time (min)
C
D
Height (nm)
30.0
10.0
-10.0
E
F
G
Height (nm)
Distance (μm)
H
Height (nm)
Distance (μm)

**Supplementary Figure 5:** Correlative AFM/QCM-D experiments for human septin hexamers and octamers on membranes. (A) QCM-D measurements for septin hexamers (50 nM, flow rate $20\,\mu L\,min^{-1}$) (arrow marked hexamers), showing the frequency shift (top) and dissipation shift (bottom). After 150 minutes (arrow marked GTA), 0.1% glutaraldehyde (GTA) was perfused for 1 min to crosslink the network, followed by perfusion with low-salt buffer (5 min) (not shown). Successful crosslinking is evident from the decrease in the dissipation shift. We simultaneously observe a frequency shift increase, which (assuming there is no release of septins) implies that the septin film becomes more compact and thinner on GTA incubation. The estimated layer thickness decreases from 11.1 ± 0.2 nm to 6.1 ± 0.1 nm. (B) Same for septin octamers (50 nM). The estimated layer thickness decreases from 14.8 ± 0.3 nm to 12.9 ± 0.3 nm. (C) AFM image of the hexamer network on the QCM-sensor from panel A. (D) AFM image of the octamer network on the QCM-sensor from panel B. (E) Zoomed-in image of septin hexamers (boxed region in panel C). (F) Zoomed-in image of septin octamers (boxed region in panel D). (G) Height profile along the blue line in the hexamer image in panel C. (H) Height profile along the blue line in the octamer image in panel F. Scale bars, 1 μm. Color bar represents the height, with 0 nm representing the bilayer surface.

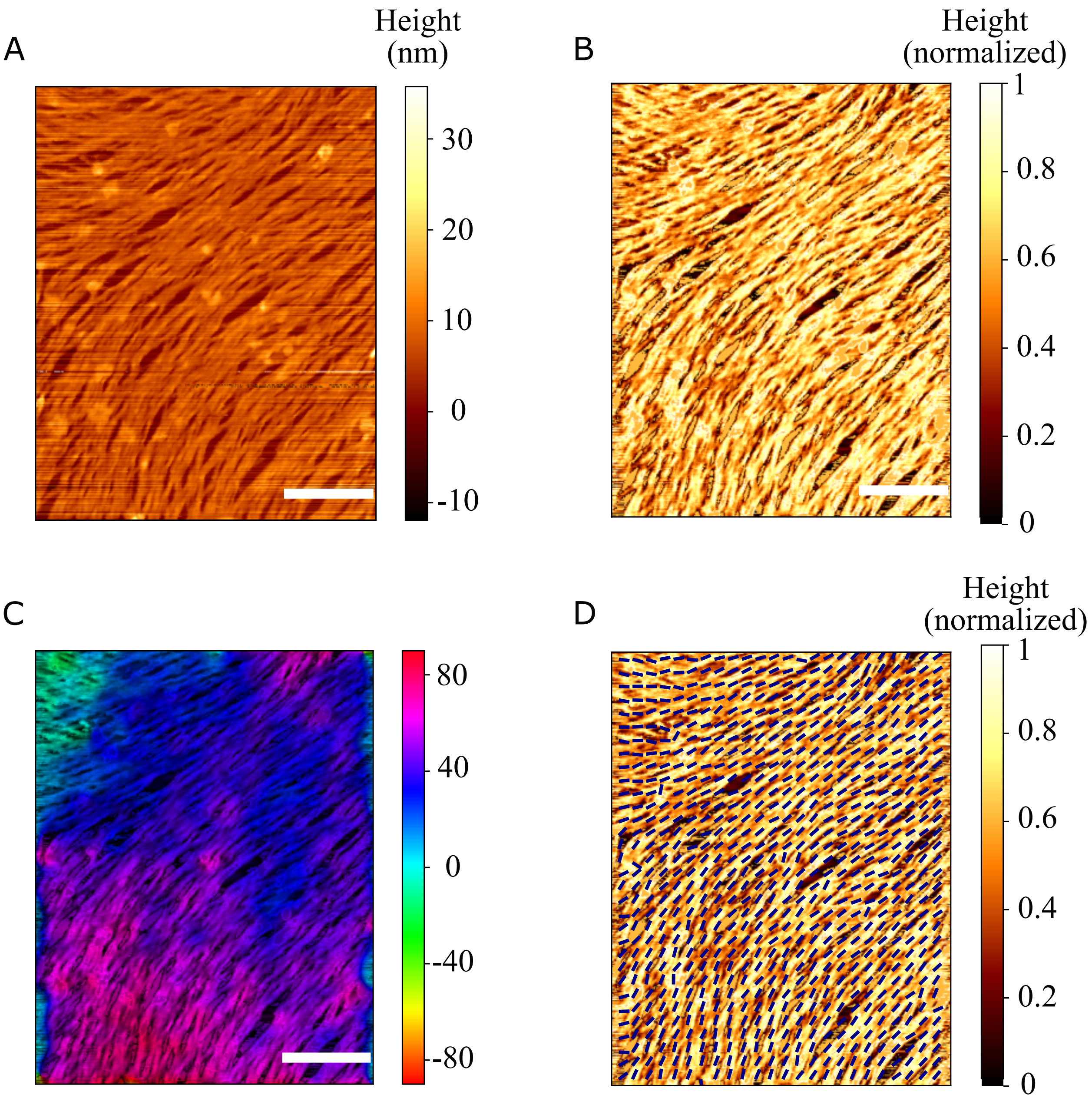


**Supplementary Figure 6:** llustration of the analysis pipeline for determining orientational order and curvature of septin filament networks on SLBs from AFM images. (A) AFM image of a septin hexamer network (100 nM) after post-processing in Gwyddion. Color bar shows height with 0 nm representing the bilayer surface. (B) Same image after filtering for line errors and height normalization using a custom Python code. Color bar shows height normalized by minimum and maximum values. (C) Overlay of processed image from panel B with color-coded pixel orientations computed with OrientationPy. Color bar shows angles in degrees (range between -90 and +90 degrees) relative to the horizontal direction. (D) Overlay of processed image from panel B with vector field (blue lines) computed with a box size of 16 by 16 pixels. Color bar shows normalized height as in panel B. Scale bars, 1 μm.

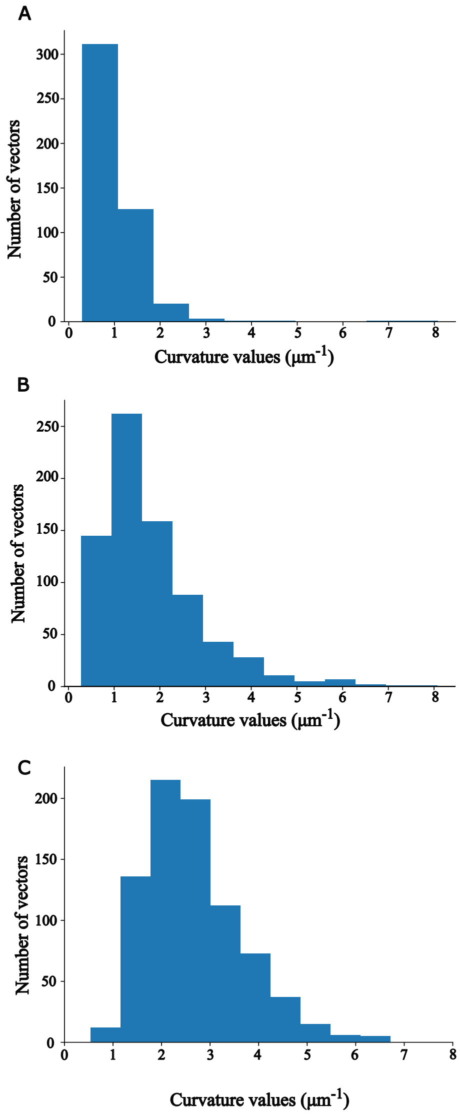


**Supplementary Figure 7:** Distribution of curvatures in 16×16 pixels regions of AFM images of septin filament networks (all 100 nM) on SLBs, calculated from the vector fields computed with OrientationPy. Examples are shown for (A) a human septin hexamer network, (B) a nematic human septin octamer network, and (C) a spiral octamer network. We observe larger curvatures for octamers than for hexamers.

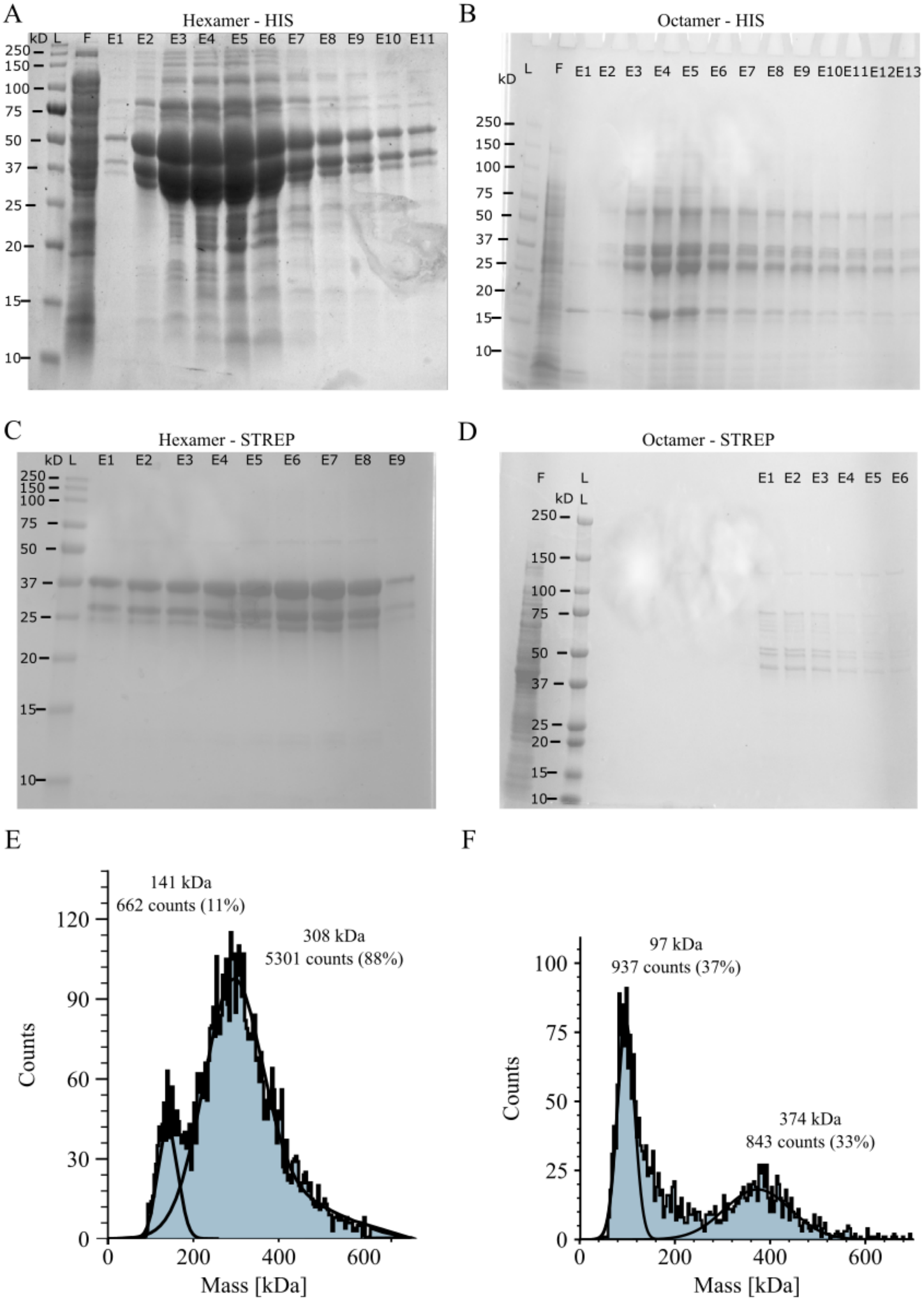
A
Hexamer - HIS
kD L F E1 E2 E3 E4 E5 E6 E7 E8 E9 E10 E11
250 150 100 75 50 37 25 20 15 10
B
Octamer - HIS
kD L F E1 E2 E3 E4 E5 E6 E7 E8 E9 E10 E11 E12 E13
250 150 100 75 50 37 25 20 15 10
C
Hexamer - STREP
kD L E1 E2 E3 E4 E5 E6 E7 E8 E9
250 150 100 75 50 37 25 20 15 10
D
Octamer - STREP
F L E1 E2 E3 E4 E5 E6
kD L
250 150 100 75 50 37 25 20 15 10
E
141 kDa
662 counts (11%)
308 kDa
5301 counts (88%)
Counts
Mass [kDa]
F
97 kDa
937 counts (37%)
374 kDa
843 counts (33%)
Counts
Mass [kDa]

**Supplementary Figure 8:** Characterization of purity and integrity of recombinant human septin hexamers and octamers. (A) SDS-PAGE gel after the first (His-tag) column purification for hexamers. (B) SDS-PAGE gel after the first (His-tag) column purification for octamers. (C) SDS-PAGE gel after the second (Strep-tag) column purification for hexamers. The three bands shown are SEPT2 (44 kDa), SEPT6 (48 kDa), and SEPT7 (52 kDa). (D) SDS-PAGE gel after the second (Strep-tag) column purification for octamers. The four bands shown are SEPT2 (44 kDa), SEPT6 (48 kDa), SEPT7 (50 kDa), and SEPT9i1 (67 kDa). Labels in panels A-D: (L) ladder, (F) flow-through, (E#) elution fractions selected from the chromatogram protein peak, at the start of the peak is described E1 with each following well/lane counted up until the end of the peak of the chromatogram. (E) Mass photometry for purified hexamers (12.5 nM) with Gaussian fits to the peaks, showing a peak for tetramers at 141 kDa and a peak for hexameric complexes at 308 kDa. (F) Mass photometry for purified octamers (12.5 nM) with Gaussian fits to the peaks, showing a peak for dimers (97 kDa) and a peak for octameric complexes (374 kDa).

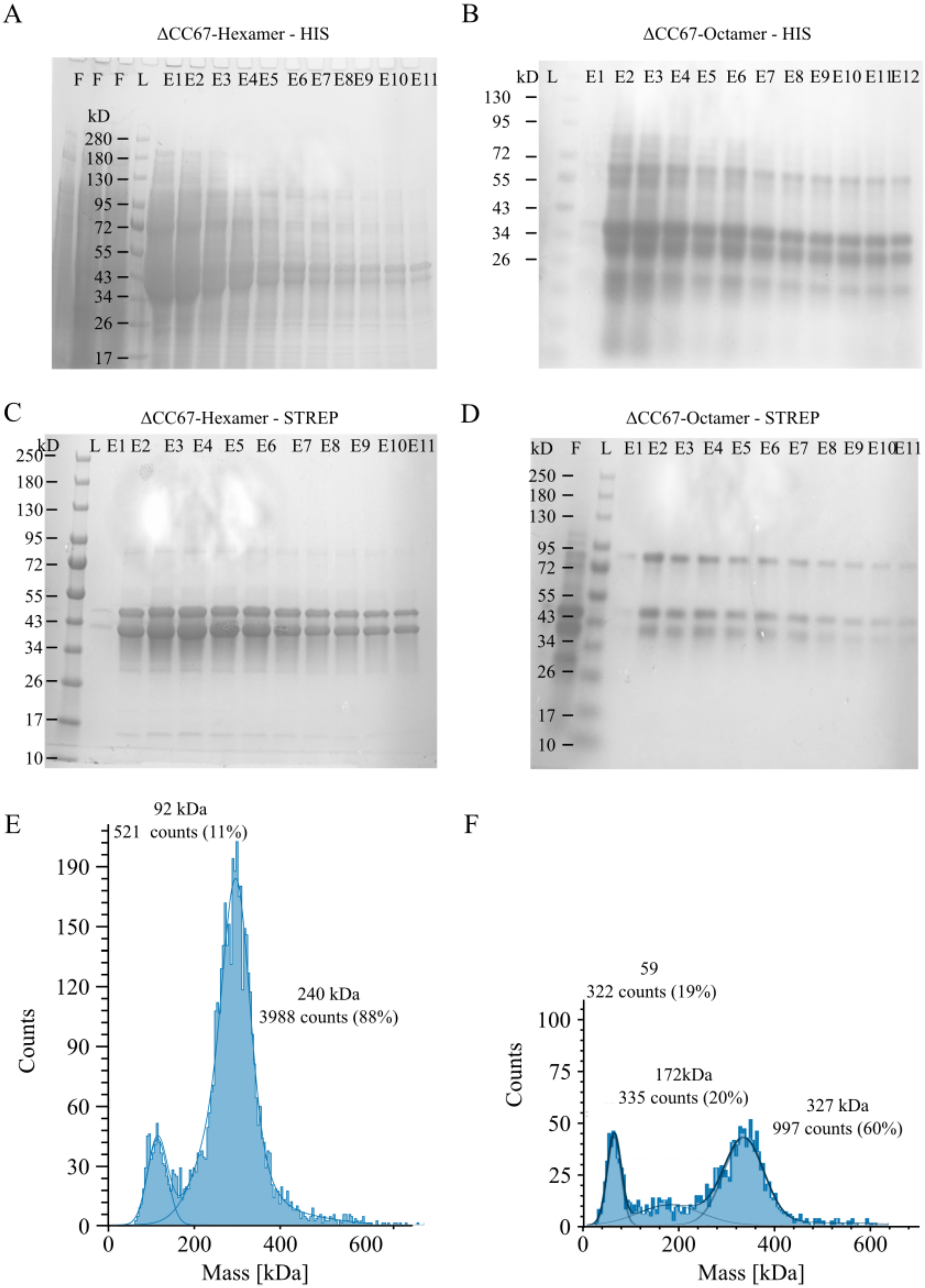
A
ΔCC67-Hexamer - HIS
F F F L E1 E2 E3 E4 E5 E6 E7 E8 E9 E10 E11
kD
280
180
130
95
72
55
43
34
26
17
B
ΔCC67-Octamer - HIS
kD L E1 E2 E3 E4 E5 E6 E7 E8 E9 E10 E11 E12
130
95
72
55
43
34
26
C
ΔCC67-Hexamer - STREP
kD L E1 E2 E3 E4 E5 E6 E7 E8 E9 E10 E11
250
180
130
95
72
55
43
34
26
17
10
D
ΔCC67-Octamer - STREP
kD F L E1 E2 E3 E4 E5 E6 E7 E8 E9 E10 E11
250
180
130
95
72
55
43
34
26
17
10
E
92 kDa
521 counts (11%)
240 kDa
3988 counts (88%)
Counts
Mass [kDa]
F
59
322 counts (19%)
172kDa
335 counts (20%)
327 kDa
997 counts (60%)
Counts
Mass [kDa]

**Supplementary Figure 9:** Characterization of purity and integrity of recombinant human septin hexamers and octamers with truncated SEPT6 and SEPT7 coiled coils (Δ67 mutants). (A) SDS-PAGE gel after the first (His-tag) column purification for ΔCC67-hexamers. (B) SDS-PAGE gel after the first (His-tag) column purification for ΔCC67-octamers. (C) SDS-PAGE gel after the second (Strep-tag) column purification for ΔCC67-hexamers. The three bands shown are SEPT2 (44 kDa), ΔCC-SEPT6 (35 kDa), and ΔCC-SEPT7 (38 kDa). (D) SDS-PAGE gel after the second (Strep-tag) column purification for ΔCC67-octamers. The four bands shown are SEPT2 (44 kDa), ΔCC-SEPT6 (35 kDa), ΔCC-SEPT7 (36 kDa), and SEPT9i1 (67 kDa). Labels in Fig A-D: (L) ladder,(F) flow-through, (E#) elution fractions selected from the chromatogram protein peak, at the start of the peak is described E1 with each following well/lane counted up until the end of the peak of the chromatogram. (E) Mass photometry for purified ΔCC67-hexamers (12.5 nM) with Gaussian fits to the peaks, showing a peak for tetramers (92 kDa) and one for hexamers (240 kDa). (F) Mass photometry for purified ΔCC67-octamers (12.5 nM) with Gaussian fits to the peaks, showing peaks for dimers (59 kDa), tetramers (172 kDa) and octamers (327 kDa).
